# FuChi: A cell cycle biosensor for investigating cell-cycle kinetics during avian development

**DOI:** 10.1101/2025.09.24.678103

**Authors:** Zoe R. Sudderick, Tiernan Briggs, Shirooza Mubarak, Melinda Van Kerckvoorde, Ana R. Hernandez-Rodriguez, Sudeepta K. Panda, Jon Riddell;, Cameron Batho-Samblas, Lorna Taylor, Lynn McTeir, Dominique Meunier, Amy Findlay, Flossie Roberts, Anna Raper, Asako Sakaue-Sawano, Atsushi Miyawaki, Joe Rainger, Jeffrey J. Schoenebeck, Cornelis J. Weijer, Mike J. McGrew, Denis J. Headon, Megan G. Davey, Richard L. Mort, James D. Glover

**Affiliations:** The Roslin Institute & R(D)SVS, University of Edinburgh, Edinburgh, UK; Roslin Institute Chicken Embryology (RICE) group; Division of Biomedical and Life Sciences, Faculty of Health and Medicine, Lancaster University, Lancaster, UK; School of Life Sciences, University of Dundee, Dundee, UK; Gurdon Institute, University of Cambridge, Cambridge, UK; Laboratory for Cell Function Dynamics, RIKEN Center for Brain Science, Wako, Saitama 351-0198, Japan; Division of Physiology and Climatology, ICAR-Indian Veterinary Research Institute, Izatnagar, 243 122, U.P., India

## Abstract

The ability to monitor the proliferative status of live cells both *in vitro* and *in vivo* over time has revolutionised our understanding of development, growth and disease. This was first made possible by fluorescent ubiquitination-based cell cycle indicator (Fucci) technology, which distinguishes specific cell cycle phases through the reciprocal degradation of fluorescently tagged, truncated forms of human CDT1 and GMNN proteins. Fucci genetic systems have been successfully implemented in transgenic mice, zebrafish, and axolotls. To date, no viable, stably expressing Fucci line has been developed in an avian species. Although a range of continuously improving Fucci constructs have been developed in recent years, existing *in vivo* Fucci models remain limited because they rely on older reporter technology that fails to distinguish cells in S, G2, and M phases or to label cells in early G1. As a result, these models can be challenging to interpret and their utility for continuous cell tracking and precise analysis of cell-cycle dynamics is limited. Here, we introduce FuChi, a multicistronic Fucci-expressing chicken line incorporating a newly optimised reporter construct composed of an mCerulean-tagged Histone H1.0 linker protein fused via a self-cleaving 2A peptide to the tandem Fucci(CA) cell cycle biosensor, with additional epitope tags included for detection in fixed tissues. We show that this system accurately discriminates and permits tracking of cells in G1, S, G2, and M phases both *in vitro* and *in vivo*, enabling faithful visualisation of cell cycle status in intact tissues and organs. Using FuChi embryos, we mapped proliferation dynamics across developing tissues, analysed cell cycle states of migrating cells, and performed live imaging of early embryos. These latter experiments revealed that transition from S phase may be a key morphogenetic event during gastrulation as mesendoderm cells egress from the primitive streak to form embryonic structures including the prechordal plate. Pairing this advanced reporter with the intrinsic experimental advantages of the chicken embryo positions FuChi as a premier *in vivo* system for studying cell-cycle kinetics in development, delivering clear technological improvements over current Fucci models. FuChi chickens provide a powerful new resource for studying embryonic development, organ growth, tissue homeostasis, disease processes, and infection responses.

## Introduction

The ability to monitor the proliferative status of cells has transformed our view of development, growth, and disease. Cell cycle control diversifies as development proceeds^1^ and is tightly coordinated within tissues, underpinning key aspects of morphogenesis and organogenesis such as kidney ureteric bud branching and digit formation^2^. In recent years, live imaging of cell cycle progression has become the gold standard for cell cycle analysis.

The E3 ubiquitin ligases APC^Cdh^^1^, SCF^Skp^^2^, and CUL4^Ddb^^1^ regulate the cell cycle by targeting specific proteins for ubiquitin-mediated degradation. SCF^Skp^^2^ acts not only as a substrate of APC^Cdh^^1^ but also directly inhibits it, resulting in reciprocal oscillations in their activities and the levels of their target proteins. APC^Cdh^^1^ is predominantly active in late mitosis and G1 phase, SCF^Skp^^2^ functions during S and G2 phases, and CUL4^Ddb^^1^ is predominantly active during S phase^3–6^. CDT1 and GMNN, which regulate DNA replication licensing, are targets of SCF^Skp^^2^/CUL4^Ddb^^1^ and APC^Cdh^^1^, respectively, and therefore also exhibit oscillations in their protein levels (Figures 1A and B)^7,8^. These dynamics have been exploited to develop the Fucci (Fluorescent Ubiquitination-based Cell Cycle Indicator) systems (Figures 1C and 1D)^9–11^.

**Figure 1.**
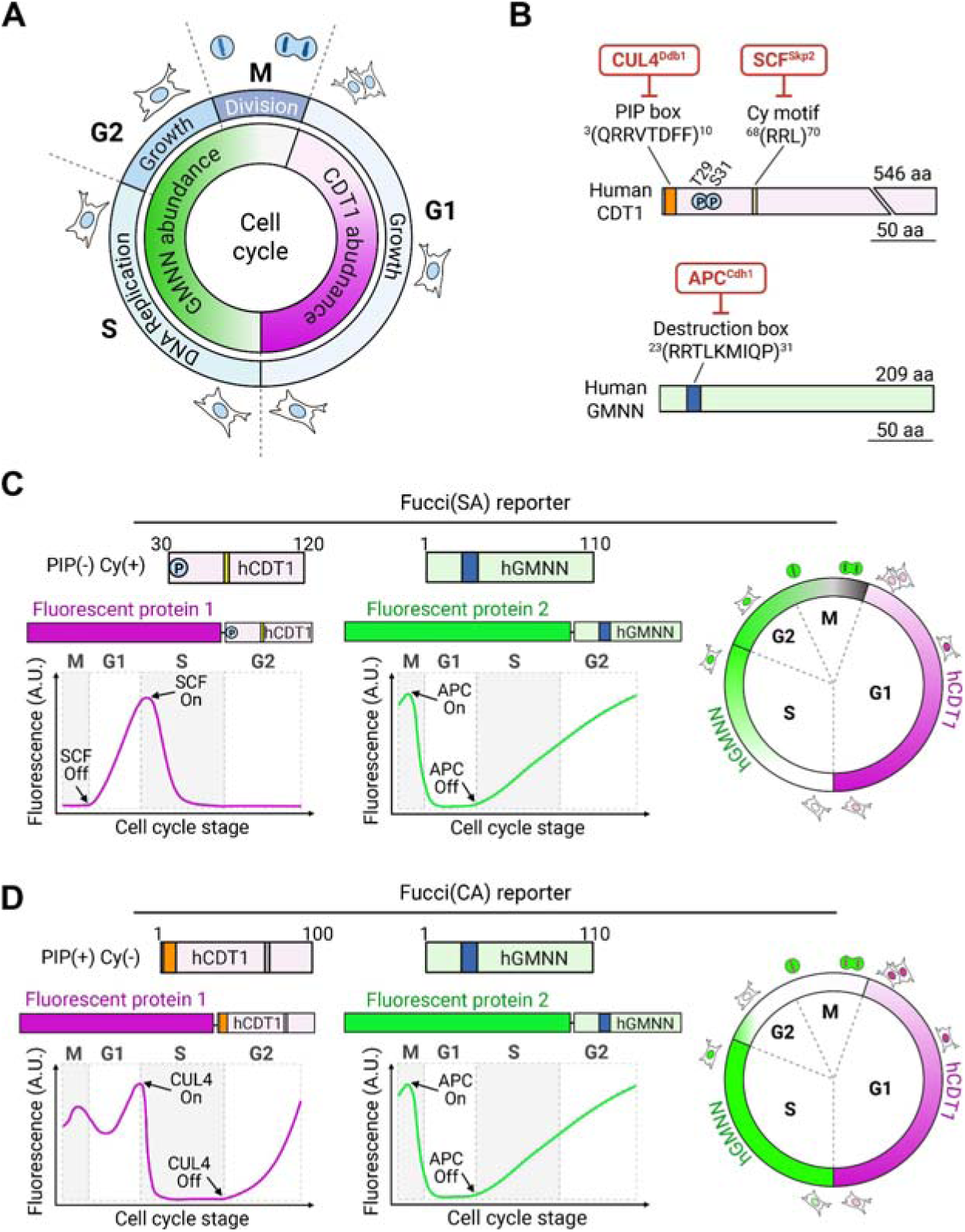
Basis and design of different Fucci reporters. **A)** Schematic of cell cycle stages including the abundance of GMNN and CDT1 proteins, two essential and interacting DNA replication licensing factors expressed at distinct phases of the cell cycle. For visualisation purposes, we have chosen to represent the red fluorescence as magenta in all our Fucci reporter system schematics. **B)** Structure of human CDT1 (hCDT1) and GMNN (hGMNN) proteins, including key features regulating their proteolysis during the cell cycle. The N terminal of CDT1 is targeted for degradation by two different E3 ubiquitin ligases, CUL4^Ddb^^1^ and SCF^Skp^^2^, which bind to the PIP box and Cy motif, respectively. Phosphorylation of key amino acids (Thr29 and Ser31) facilitate binding of SCF^Skp^^2^. Ubiquitination and proteolysis of GMNN is regulated through the E3 ligase APC^Cdh^^1^ binding to the destruction box. **C)** Fucci biosensors utilise fusion proteins of hCDT1 and hGMNN, containing truncated forms of the proteins fused with fluorescent markers. In Fucci(SA) systems, an SCF^SKP^^2^-sensitive human CDT1 probe linked to a fluorescent protein (e.g. mKO2 / mCherry) is used in combination with a truncated human APC^Cdh^^1^-sensitive GMNN1 fused to a spectrally distinct fluorescent protein (e.g. mAG / mVenus). The expected fluorescent profiles throughout the cell cycle reflect hCDT1 and hGMNN1 degradation by their respective E3 ligases. Note that double labeled cells are white and negative cells are grey in the combined schematic. **D)** Fucci(CA) systems combines CUL4^Ddb^^1^-sensitive and APC^Cdh^^1^-sensitive GMNN1 probes, which creates a distinct fluorescent profile from Fucci(SA) based on the degradation of hCDT1. The grey Cy motif denotes the RRL to AAA change in this probe, which renders the Cy motif nonfunctional. As above, double labeled cells are white and negative cells are grey in the combined schematic.

The original Fucci(SA) probe pair (Figure 1C) included a fusion of monomeric Kusabira Orange (mKO2) with a truncated human CDT1 (amino acids 30–120), and a fusion of monomeric Azami Green (mAG) with the N-terminal amino acids (1-110) of human GMNN. The mKO2-hCDT1(30/120) probe accumulates in G1 phase, labelling cell nuclei red, and is degraded at the G1–S transition, while the mAG-hGem(1/110) probe accumulates during S, G2, and M phases, labelling cell nuclei green, and is rapidly degraded before cytokinesis^11^ (Figure 1C).

An improved version of Fucci(SA), sometimes referred to as Fucci2 or Fucci(SA)2, replaces the original mKO2 and mAG fluorescent proteins with mCherry and mVenus, respectively, increasing signal separation and compatibility with other reporters^12^. This probe set was initially expressed from separate constructs but incorporation of a 2A self-cleaving peptide enabled equimolar expression of reporter proteins from a single promoter (termed Fucci2a or tandem-Fucci(SA)2) and facilitated the development of Cre-inducible mouse models^10,13^.

The bicistronic tandem-Fucci(CA) probe pair^9^ (Figure 1D) updates the CDT1 probe from Fucci(SA) by removing the SCF^Skp^^2^-targeted Cy motif and instead including an N-terminal PIP box, which is targeted for degradation in S-phase via CUL4^Ddb^^1^. This leads to accumulation of CDT1 starting at the S-G2 transition and continuing through G2, M and G1 phases with rapid degradation at the G1/S transition. This results in labelling of G1-phase nuclei as red, S-phase nuclei as green and G2 and M phase nuclei as yellow with just two sensors^9^ (Figure 1D).

Fucci(SA) has primarily been used in mouse and human cell lines^11,13–16^ while tandem-Fucci(SA) has also been deployed in mouse models^10,14,15^ and has been optimised for use in zebrafish, xenopus (zFucci)^17,18^ and in axolotls^19^ using species-specific homologues of Cdt1 and Geminin. However, none of these *in vivo* models label all four cell cycle phases, and detecting the fluorescent proteins in fixed tissues remains challenging. Fucci4 solves the first of these problems by combining four sensors: hCDT1(30-120) degraded by SCF^Skp^^2^ marks G1, a PIP degron targeted by CUL4^Ddb^^1^ labels S phase, GMNN(1–110), degraded by APC^Cdh^^1^, marks S and G2, and an H1.0 linker histone fusion highlights chromatin during mitosis. However, although powerful for *in vitro* imaging Fucci4 is not easily deployable in animal models as the sensors are expressed separately^15^.

Chickens are a valuable research model because their embryos develop externally, allowing real-time imaging and manipulation, and their early amniote morphology closely parallels that of humans^20,21^. Fucci systems have previously been introduced into cultured chicken fibroblasts and primordial germ cells, respectively^22,23^, suggesting compatibility with avian species; however, no tractable *in vivo* germline model system has yet been developed.

Here, we describe a new Fucci-based biosensor, H1.0-Fucci(CA)2, designed to label all four cell cycle phases and incorporating epitope tags for detection in fixed tissues. Using the PiggyBac system, we integrated H1.0-Fucci(CA)2 into chicken fibroblasts and primordial germ cells, and demonstrate that it outperforms Fucci(SA)2 by faithfully discriminating between cells in G1, S, G2, and M phases. We further generated a stable line of H1.0-Fucci(CA)2-expressing transgenic chickens (FuChi), enabling, for the first time, *in vivo* studies of cell cycle progression during time lapse embryonic development in an avian model.

## Materials and Methods

### H1.0-Fucci(CA)2 construction

To generate pcDNA5-CAG-H1.0Cerulean-Fucci(CA), a gene fragment encoding the *mCerulean* fluorescent protein fused to an HA epitope tag and a porcine teschovirus-1 2A peptide (P2A), flanked by 5′ MluI and 3′ HindIII restriction sites, was synthesised by GeneArt (Thermo Fisher). The *mCerulean* fragment was cloned into pZERO-1 (Thermo Fisher), and the coding sequence of the mouse *H1f0* gene (encoding the H1.0 linker histone) was PCR-amplified from mouse cDNA (Table 1) and cloned into the MluI site.

**Table 1:**
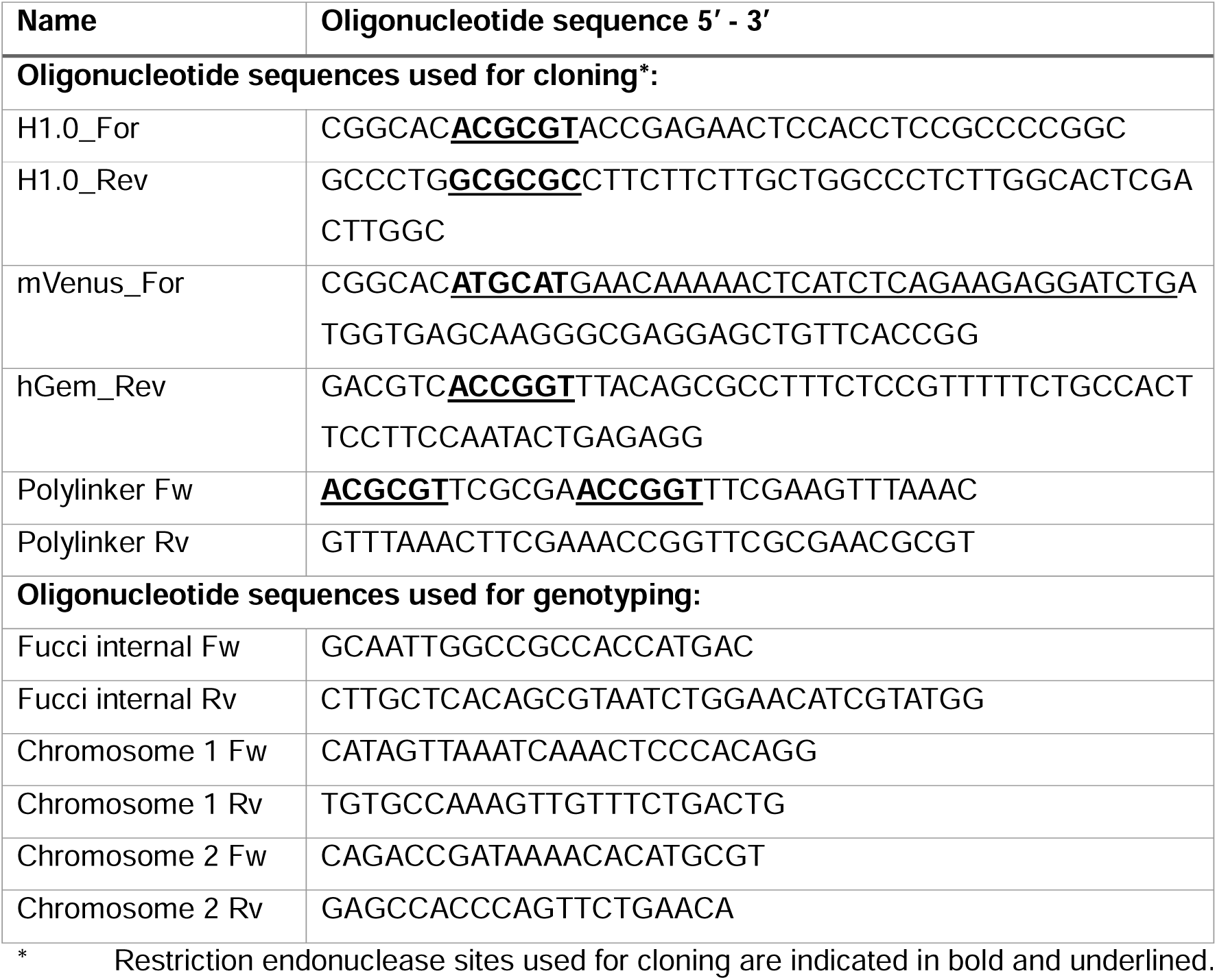
Oligonucleotide sequences used for cloning and PCR.

Subsequently, an *mCherry-hCDT(1/100)Cy(-)* gene fragment^9^,including a 5′ 6xHis tag and HindIII, and 3′ NsiI sites, was synthesised by GeneArt. An *mVenus-hGem(1/110)* fragment^9^ was PCR-amplified to incorporate a 5′ Myc tag, NsiI, and 3′ AgeI sites (Table 1). These three components were sequentially cloned into a pCDNA5-Flp backbone modified to contain the CAG promoter in place of CMV^13^. The fidelity of all clones was checked by sequencing.

To generate pB-H1.0Cerulean-Fucci(CA)2, a polylinker sequence (Table 1) was cloned into pB-H2BCerulean-Fucci2a^24^ using MluI/PmeI, and the H1.0-Fucci(CA) construct was transferred as an MluI/AgeI fragment.

### Transfection into PGCs

To generate H1.0-Fucci(CA)2 PGCs, 2 µg of pB-H1.0Cerulean-Fucci(CA)2 was co-transfected with 1 µg of piggyBac Hybase vector^25^ into a previously derived line of male PGCs (FR2M, Hyline birds) using Lipofectamine 2000 (Life Technologies), and grown in FAOT medium^26^. Stably transduced cells were then selected with puromycin (Sigma Aldrich) and expanded prior to PGC injection.

### Generation of FuChi embryos

H1.0-Fucci(CA)2 PGCs were micro-injected into HH15-16 surrogate iCaspase9 sterile hosts as described previously^27^. Briefly, 1□µl of 25□mM B/B compound (Takara Bio) was added to a 50□µl cell suspension (5000 PGCs/µl) before 1 µl of this suspension was injected into the dorsal aorta of host embryos within windowed eggs. Egg shells were sealed with medical Leukosilk tape (BSN Medical) and incubated until hatch. Founder F_0_ FuChi males were mated with wild type Hyline females in pens to produce heterozygous F_1_ FuChi eggs and offspring. Chicken experiments were conducted under UK Home Office licence PP9565661 and approved by the Roslin Institute Animal Welfare and Ethical Review Board Committee. The fertility, viability (of fertile eggs), and transgene transmission rates (determined by fluorescence) were recorded.

### Eggs

The FuChi line is maintained in the National Avian Research Facility (NARF), Roslin Institute, UK. Eggs were incubated vertically in rocking or non-rocking incubators (Masalles) at 37.8°C. All post incubation day 14 embryos were treated in accordance with the ASPA 1986 schedule 1 culling methods. Fluorescent, and therefore transgenic, embryos were identified using a Nightsea SFA-RB fluorescence adapter and/or a Zeiss Axiozoom V6 microscope.

### Cell culture and transfections

DF1 cells were maintained in DMEM (Life Technologies) containing 10% FBS, 2% chicken serum (CS), 1% penicillin/streptomycin. Cells were transfected with either pB-H1.0Cerulean-Fucci(CA)2 or pB-H2BCerulean-Fucci2A^24^, together with a vector expressing piggyBac Hybase using Lipofectamine LTX (Life Technologies) as per the manufacturer’s instructions. The cell lines were termed H1.0-Fucci(CA)2-DF1 and H2B-Fucci(SA)2-DF1 in line with Sakaue-Sawano et al.^9^.

FuChi primary fibroblasts were derived from HH19 limb bud micromasses. HH19 embryos were dissected from the egg into DMEM and all limb buds removed. Limb buds were placed in 10X Trypsin (Life Technologies) on ice for 30-40 minutes, followed by two washes with PBS and removal of the ectoderm with sharpened tungsten needles. Limb bud mesoderm was collected in 100 μl of micromass culture medium (50:50 DMEM/Ham’s F12, 10% FCS, 1% Antibiotic/Antimycotic, 1% L-Glutamine) and dissociated by trituration. 900 μl of cell culture medium was added and the samples were centrifuged at 300 rcf for 2 minutes. The cell pellet was resuspended in 10 μl of cell culture media per limb bud. 10 μl drops were added to individual wells of a 24 well plate and incubated for 30 minutes to 1 hour at 37°C. Wells were then flooded with 1 ml of micromass culture media

To derive FuChi PGCs, 1-2 µl of blood was extracted from HH16 F_1_ FuChi embryos, transferred into FAOT medium and cells expanded as previously described^26^.

### Cell treatments

Prior to flow cytometry analysis, H2B-Fucci(SA)2-DF1, H1.0-Fucci(CA)2-DF1 cells or FuChi PGCs were grown in their regular medium conditions as described above. Cells were then treated for 16 hours in control (regular medium plus DMSO), base (regular culture medium minus FBS and CS for DF1, or removal of FAOT growth factors for PGCs) or Nocodazole treated (regular culture media containing 100 ng/ml Nocodazole (Sigma Aldrich) conditions. Following treatment, cells were dissociated from the flasks / PGCs removed, centrifuged, put through a 70 µm cell strainer, and stained with DAPI (Sigma Aldrich) for 30 minutes. Cells were analysed for DNA content and mCherry/mVenus fluorescence using a BD Fortessa X20.

### Live cell imaging

Primary FuChi limb micromass cultures were grown on glass bottom culture plates in micromass culture medium and imaged for 72 hours. H2B-Fucci(SA)2-DF1 and H1.0-Fucci(CA)2-DF1 cultures were grown on glass bottom plates in cell culture media overnight then imaged for 48 hours. All live imaging was carried out on a Zeiss LSM880 confocal microscope, with z-stack images taken at 10-minute intervals. Cell tracking and generation of individual cell montages was carried out using ImageJ/Fiji and the Fucci Tools plugin^28^.

### Lightsheet imaging and analysis

FuChi eggs were obtained from the NARF and stored at 16°C until the start of the experiments. The eggs were incubated for 1 hr to equilibrate to 37°C, after which embryos were isolated on filter paper and mounted in a specially constructed imaging chamber and imaged on a dedicated lightsheet microscope as described previously^29^. To image the embryos at 10x magnification two slightly overlapping scans were performed. During the scanning the surface of the embryo was kept in focus using a dedicated surface tracking algorithm as previously described^29^. Each scan consisted of 2500-3000 successive slices taken at 1.84 µm intervals and imaged at 488 and 562 nm excitation. This procedure allowed us to collect whole embryo images at two wavelengths at 8-10 minute intervals.

To process the raw image data, the images were transformed into a rectangular coordinate system and the surface was detected using a surface finding algorithm. The top 30 z slices (14 µm) were projected onto a plane and the nuclei detected in each channel using a custom Matlab Laplacian of Gaussian (LoG) blob detector algorithm. Nuclei were classified into three classes, those detected exclusively in either the green or red channel and those detected in both, as determined by their co-localisation. For analysis of variation of hGMNN-mVenus (488 channel) and hCDT1-mCherry (562 channel) expression during the cell cycle *in vivo*, selected nuclei were traced manually and their intensities in both channels measured. Individual traces were aligned using the sudden breakdown of hGMNN-mVenus and hCDT1-mCherry at mitosis. The boundary between the embryonic and extra embryonic area at the start of the experiment was derived from the Lagrangian repeller^30,31^ and subsequently evolved for all successive timepoints using calculated tissue velocity^29^.

### Whole genome sequencing

Sequencing was carried out as in Oh et al.^32^. Briefly, E9 livers were snap frozen on dry ice and stored at -80°C. Genomic DNA was extracted from the frozen samples using an Monarch HMW DNA Extraction Kit (New England Biolabs) per manufacturer’s instructions. Diluted genomic DNA quality was assessed using an Agilent TapeStation, and concentrations were quantified with a Qubit fluorometer. gDNA from 5 samples was pooled. Library preparation and sequencing was carried out by Edinburgh Genomics. The SMRTbell library was sequenced on a Pacific Biosystems Revio platform (25M ZMW, HiFi mode) using a single SMRT cell. A total of 58 Gb of DNA sequence was generated, with an average fragment size of 19 kb and a mean Phred quality score of Q30. Data was returned in bam and fastq file formats (ENA accession: PRJEB98847). To detect the genomic location(s) of transgene insertion, hi-fi reads in fastq.gz format were first aligned against the corresponding transgene sequence (using Minimap2 (v2.24-r1122)^33^. In total, 69 of 2,983,364 hi-fi reads were mapped. Transgene-mapped reads were retained by Samtools (v1.18 using htslib v1.18)^34^. The output file was converted from bam to fastq and the transgene-mapped reads were realigned using Minimap2 against the paternal haplotype of GRCg7w (white leghorn layer, GCF_016700215.2)^35^.

### PCR

Insertion sites in chromosomes 1 and 2 revealed by long read sequencing were confirmed by PCR (See Table 1 for primers) using Phusion polymerase (NEB), and the supplied HF or GC buffers. Detection of transgene insertion on Chromosome 1 was carried out with HF buffer with the following conditions: 98°C/30s, (98°C/10 s, 60°C/30 s, 72°C/30 s) x33, 72°C/5 mins. Detection of transgene insertion on Chromosome 2 was carried out with GC rich buffer with the following conditions: 98°C/30s, (98°C/10 s, 64°C/30 s, 72°C/30 s) x30, 72°C/5 mins. PCR samples were run on 2% agarose gels for assessment.

### Protein sequence comparison

Cdt1 and Gmnn protein sequences from various species were downloaded from NCBI and aligned in MegAlign Pro 17 (DNAstar) using the MAFFT algorithm.

### Cryo-embedding and paraffin processing

For cryo-processing, FuChi embryos were dissected from the egg, briefly washed in PBS, and fixed with 4% PFA or 10% NBF. Following fixation, embryos were washed in PBS then equilibrated in 15% sucrose / PBS at room temperature, then incubated in 15% sucrose / 7.5% gelatin / PBS at 37°C until fully equilibrated. Blocks of samples embedded in sucrose/gelatin blocks were flash frozen in isopentane on dry ice, and stored at -80°C. 10 μm cryosections were cut on a Leica CM1950 Cryostat. For imaging, cryosections were incubated in PBS at 37°C for 15 minutes to remove gelatin, before mounting in Prolong Gold (Life Technologies) and imaged on a Zeiss LSM880 confocal microscope.

For paraffin embedding, fixed samples (10% NBF or 4% PFA) were washed in PBS, dehydrated through 70%, 90%, and 100% ethanol before clearing in 1:1 ethanol: Chloroform, and a final wash in 100% Chloroform. Samples were incubated overnight in paraffin wax at 65°C prior to embedding. 6 μm sections for immunofluorescent analysis were cut using a Thermo Microtome.

### Immunofluorescence and fluorescent RNA in situ hybridisation

Immunofluorescent detection of the epitope probes in the H1.0-Fucci(CA)2 biosensor was carried out on paraffin and cryo- sections of FuChi embryos using anti-6x-His Tag (Mouse (1:100), Thermo Fisher #MA1-21315) and anti-Myc-Tag (Rabbit (1:100), Cell Signaling Technologies #71D10) antibodies. For paraffin sections, after dewaxing, an antigen retrieval step was performed with Tris-EDTA using an Antigen Retriever 2100 (Aptum Biologics). For cryosections, samples were equilibrated to room temperature and then incubated for 15 minutes in PBS at 37°C to remove gelatin. After these steps, paraffin and cryo- sections were treated similarly. Sections were permeabilised with PBS containing 0.5% Triton-X100, briefly washed in PBS containing 0.1% Tween 20 (PBST), and blocked in 5% goat serum / PBST for one hour at room temperature. Sections were then incubated overnight at 4°C in primary antibodies (anti-6x His Tag, anti-Myc Tag, anti-YAP1 (Rabbit (1:200) Proteintech #13584) diluted in blocking buffer. Samples were washed in PBST before incubation in fluorescent secondary antibodies (Goat anti-rabbit IgG Alexafluor 647 (1:500, Life Technologies # A-21245), Goat anti-mouse IgG Alexafluor 488 (1:500, Life Technologies, #A-11001) diluted in blocking buffer for one hour at room temperature. Samples were washed in PBST, counterstained with DAPI, mounted in Prolong Gold and imaged on a Zeiss LSM880 confocal microscope.

For wholemount immunofluorescence, FuChi embryos were dissected and fixed in 10% NBF. Samples were washed 3 x 30-60 mins in PBS containing 0.5% Triton X-100 (PBTx) with rocking, before blocking in 10% goat serum / PBTX for 2 hrs at room temperature with rocking. Samples were then incubated with anti-DAZL antibody (Rabbit (1:250), Abcam, #AB215718) diluted in blocking buffer for 20 hrs at 4°C with rocking. Following primary antibody incubation, samples were washed 5 x 30-60 mins in PBTx, before incubation with secondary antibody (Goat anti-rabbit IgG Alexafluor 647;1:200) for 20 hours at 4°C with rocking. Samples were then washed 5 x 30-60 mins in PBTx, counterstained with DAPI (1:2000 in PBTX) for 1 hour, mounted in Prolong Gold and imaged on a Zeiss LSM880 confocal microscope.

Fluorescent *in situ* hybridisation was performed by hybridisation chain reaction (HCR; Molecular Instruments). Briefly, FuChi embryos were dissected, and immediately fixed in 4% PFA at room temperature for 1 hour protected from light. Samples were rapidly dehydrated (5 mins per dilution series) into 100% methanol, stored at -20°C for 1-2 hrs, before rehydration. HCR was then performed on samples following the manufacturer’s standard protocol using *PTCHD1* and *SHH* probes designed and ordered directly from Molecular Instruments. After HCR, embryos underwent wholemount imaging or were processed for cryosectioning and sections imaged using a Zeiss LSM880 confocal microscope.

### Early embryo culture and imaging

Early embryo cultures were performed as in Chapman et al.^36^ Briefly, embryos were collected on filter paper, washed in Ringer’s solution to remove excess yolk, before culturing ventral side up on glass bottomed six well plate containing a thin layer of albumin / agar (0.3%) mix. Embryos were incubated in a 37°C imaging chamber, with wet tissue included in the surrounding space and/or unused wells on the plate to ensure humidity, and imaged on a Zeiss LSM880 confocal microscope.

### Fluorescence quality assessment

E3 embryos were imaged *in ovo*, immediately upon removal, and at 4, 24, 48, 72, and 168 hrs. All images were taken with the same fluorescence settings on a Zeiss Axiozoom V16 microscope. Samples were fixed in either 4% PFA or10% NBF for 2, 4, or 24 hours, then stored in PBS or 70% ethanol.

### Image based determination of cell cycle status in PGCs

Following flow cytometry of FuChi derived PGC lines under different conditions, a sample of PGCs from each line and treatment group were mounted on slides and imaged using a Zeiss LSM880 confocal microscope. Using the Fucci Tools plugin for ImageJ/Fiji^28^, values of fluorescent intensity were obtained from cultured PGCs (3 independently derived lines; 3 culture conditions), *in vivo* PGCs in the germinal crescent (4 embryos), and from neighbouring somatic cells in embryo wholemounts.

Values for all cells were plotted as scatter graphs, using normalised values for mCherry and mVenus which were calculated by subtracting the population background value (mean of lowest 5 values) from the individual value and dividing this by the maximum population value mean of top 5 values). Using the percentages of PGCs in each cell cycle phase as indicated by flow cytometry analysis we determined boundaries of G1 (mCherry >0.1, mVenus <0.1), S (mCherry <0.1, mVenus >0.1), and G2/M (mCherry >0.1, mVenus >0.1). These boundaries were then applied to the values obtained from HH6 germinal crescent PGCs and surrounding somatic cells.

## Results

### Design of the H1.0-Fucci(CA)2 biosensor

To generate a Fucci-expressing chicken line that overcomes the limitations of previously described Fucci systems, we first aimed to design an optimal Fucci biosensor for chicken. Fucci(SA) sensors (Figures 1B, 1C, 2A, and 2B), the most commonly used Fucci pair, incorporate a truncated form of human CDT1(30–120) that undergoes degradation via the SCF^Skp^^2^ pathway through its Cy motif. While this system functions in mammalian models^10,11,16,37^, the Cy motif is not well conserved in non-mammalian species (Figure S1A) and does not function in zebrafish^38^, suggesting that it may not function efficiently in chicken. By contrast, Fucci(CA) systems (Figures 1B, 1D, 2A, and 2B), use CDT1(1–110) with an alternative N-terminal PIP box, which is targeted for degradation via the CUL4^Ddb^^1^ pathway and is well conserved across species (Figure S1A). The regulation of GMNN (Geminin), the counterpart in both Fucci pairs, is also highly conserved^38^ (Figure S1A). This conservation, combined with the additional advantage of S-phase discrimination offered by the PIP box–based CDT1 sensor, suggested that a Fucci(CA) pair would be the best choice for avian species.

Neither Fucci(CA) nor Fucci(SA) discriminate between G2 and M phases and the existing animal models (which are based on Fucci(SA)2) are unable to track cells through early G1^39^. A tricistronic construct, H2B-Fucci(SA)2, has been described which enabled M-phase discrimination and tracking through early G1 by labelling DNA with an H2B-Cerulean fusion protein^24^. Similarly, the Fucci4 system, although composed of individual constructs and so not useful here, used a fusion with the H1.0 linker histone to achieve the same effect^15^. However, toxicity associated with H2B fusions has also been reported *in vivo*^40^. Therefore, we selected the H1.0 linker histone fused to mCerulean (optimally spectrally separated from mVenus and mCherry) for use in our chick model. As a final optimisation step, we added HA, 6xHis and Myc tags to the H1.0-Cerulean, mCherry-CDT1(1/100)Cy(-) and mVenus-hGMNN(1/110) components, respectively, to aid in antibody detection in fixed and cleared tissues. Figure S1B shows our optimised tricistronic H1.0-Fucci(CA)2 biosensor.

### Faithful discrimination of G1, S, G2 and M phases in chicken cells

We compared the utility of our new H1.0-Fucci(CA)2 biosensor against the previously described H2B-Fucci(SA)2 biosensor^24^ by using the PiggyBac system to stably integrate each of the biosensors (Figure S1B) into immortalised chicken DF1 fibroblasts. We found that H1.0-Fucci(CA)2 accurately discriminated cell cycle stage, with similar degradation profiles to Fucci(CA)2 mammalian models^9^. This achieved a clear discrimination between G1, S and G2/M populations and a faithful response to cell cycle perturbations (Figures 2C-F, S1D, and S1F; Supplementary video 1). However, with H2B-Fucci(SA)2 we identified an extended hCDT1 degradation curve, resulting in persistence of mCherry beyond the G1/S transition (Figures 2G-J, S1E, and S1F; Supplementary video 2), making cell cycle phase difficult to determine accurately. Both the H2B-Cerulean and H1.0-Cerulean fusions resulted in bright nuclear labelling throughout the cell cycle, including early G1, with sharp peaks in fluorescence intensity as the chromosomes condense in M-phase (Figures 2C, 2E, 2G, and 2I). These results establish H1.0-Fucci(CA)2 as a superior cell cycle probe capable of functionally discriminating all cell cycle phases in chicken cells.

**Figure 2.**
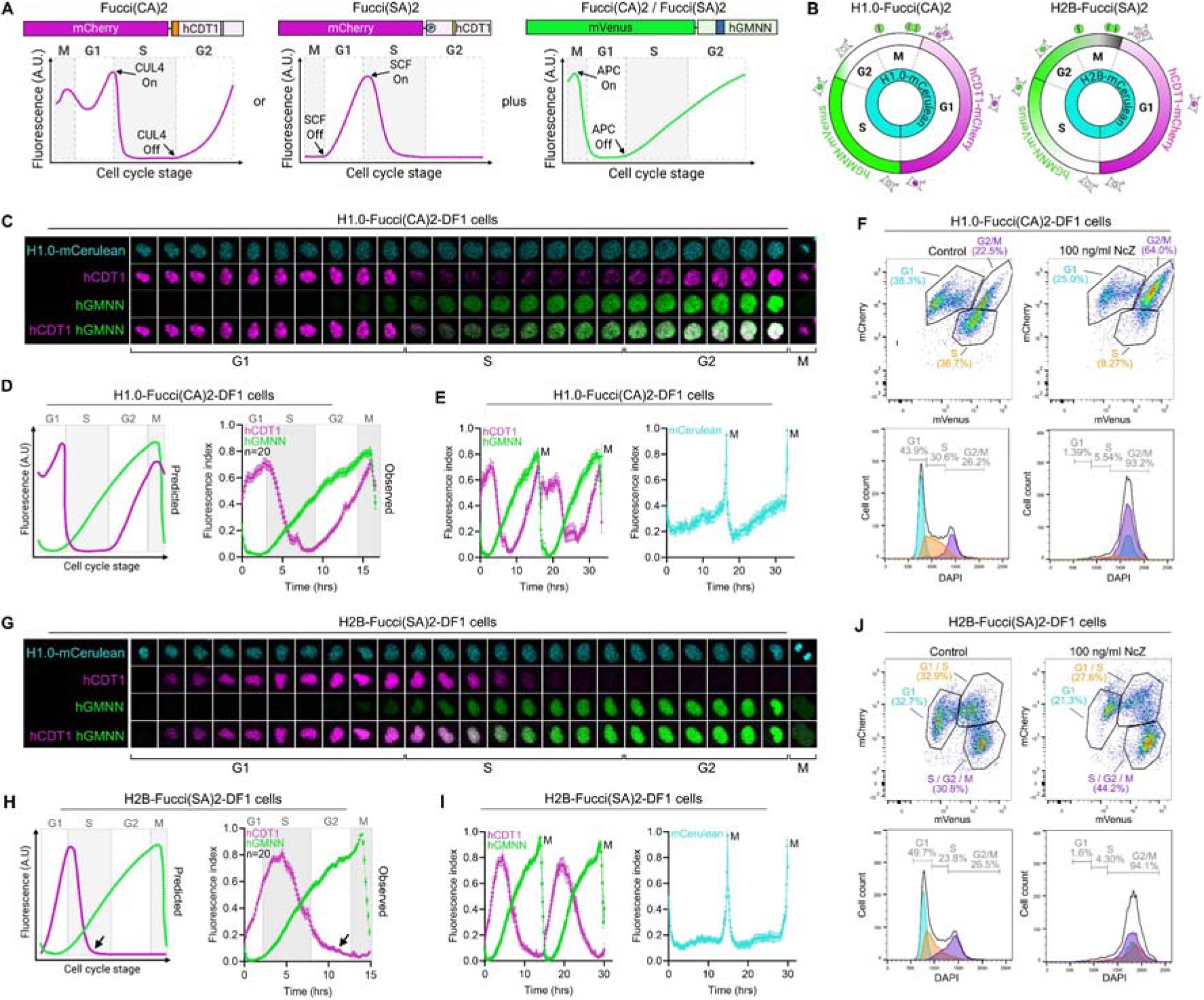
Faithful labelling of G1, S, G2 and M using H1.0-Fucci(CA)2 in chicken cells. **A)** Comparison of the mCherry-hCDT1 fusion proteins used in the Fucci(CA)2 and Fucci(SA)2 systems. The mVenus-hGMNN construct is used in both systems. **B)** Schematic showing the expected fluorescence profiles of the two systems throughout the cell cycle. The Fucci(CA)2 and Fucci(SA)2 constructs also express nuclear H1.0-mCerulean or H2B-mCerulean proteins, respectively, which are present throughout the cell cycle. Note that double labeled cells are white, negative cells are grey. **C)** Montage following an individual cell and one of its daughters through a complete cell cycle from live cell imaging of stably transduced chicken fibroblasts (DF1) expressing the H1.0-Fucci(CA)2 biosensor. **D)** Normalised fluorescence intensity plots of H1.0-Fucci(CA)2 DF1 cells throughout the cell cycle based on 20 cell division curves interpolated so that max time represents a full cell cycle. **E)** Temporal normalised fluorescence intensity profiles of H1.0-Fucci(CA)2 DF1 cells across two cell cycles based on live cell imaging data. N = 20 cells. **F)** Flow cytometric analysis of H1.0-Fucci(CA)2 DF1 cells in both control conditions and following treatment with nocodazole (NcZ) which leads to G2 arrest. Fucci(CA)2 shows faithful correlation between Fucci biosensor fluorescence and DNA content as a proxy for cell cycle stage. **G)** Montage following an individual cell and one of its daughters through a complete cell cycle from live cell imaging of stably transduced chicken fibroblasts (DF1) expressing the H2B-Fucci(SA)2 biosensor. **H)** Normalised fluorescence intensity plots of H2B-Fucci(2A)2 DF1 cells throughout the cell cycle. The observed degradation of hCDT1 is delayed in these cells compared with the expected profile (arrows). **I)** Temporal normalised fluorescence intensity profiles of H2B-Fucci(SA)2 DF1 cells across two cell cycles based on live cell imaging data. N = 20 cells. **J)** Flow cytometric analysis of H2B-Fucci(SA)2 DF1 cells in both control conditions and following NcZ. Fucci(SA)2 shows less faithful correlation between Fucci biosensor fluorescence and DNA content as a proxy for cell cycle stage in comparison to Fucci(CA)2.

### H1.0-Fucci(CA)2 expressing “FuChi” chickens

We used the piggyBac system^41^ to generate stably integrated transgenic H1.0-Fucci(CA)2 cell-cycle biosensor expressing (FuChi – “**Fu**cci **Chi**cken”) PGCs (Figure 3A). After transplantation of FuChi PGCs into sterile recipient embryos^27^, founder F_0_ males were bred with wild-type Hyline hens to generate heterozygote F_1_ embryos for analysis (Figures 3A and 3B). FuChi chickens demonstrated fertility, viability and transgene transmission rates of 88%, 93% and 49%, respectively (Figure 3C). FuChi embryos demonstrated strong H1.0-Fucci(CA)2 transgene expression with no observed lethality across all developmental stages (Figures 3B and S2A), were comparable to non-transgenic embryos (Figure 3D), could be successfully hatched (Figure S2B) and are fertile. Of note, hatched FuChi chicks are initially smaller than wild type counterparts but do grow at comparable rates (Figure S2C). However, some adult FuChi birds showed poor body condition and required euthanisia, suggesting adverse effects of the H1.0-Fucci(CA)2 transgene and/or consequences of random reporter integration via piggyBac transposition. Mapping H1.0-Fucci(CA)2 integration sites in FuChi embryos showed that most carried a single insertion on chromosome 1 (NC_052573.1:139,895,885–139,918,586) or chromosome 2 (NC_052574.1:57,700,736–57,701,209) although additional insertions exist but were not detected in the sequenced embryo pool (Figures S2D and S2E). FuChi reporter expression was consistent between embryos regardless of insertion site.

**Figure 3.**
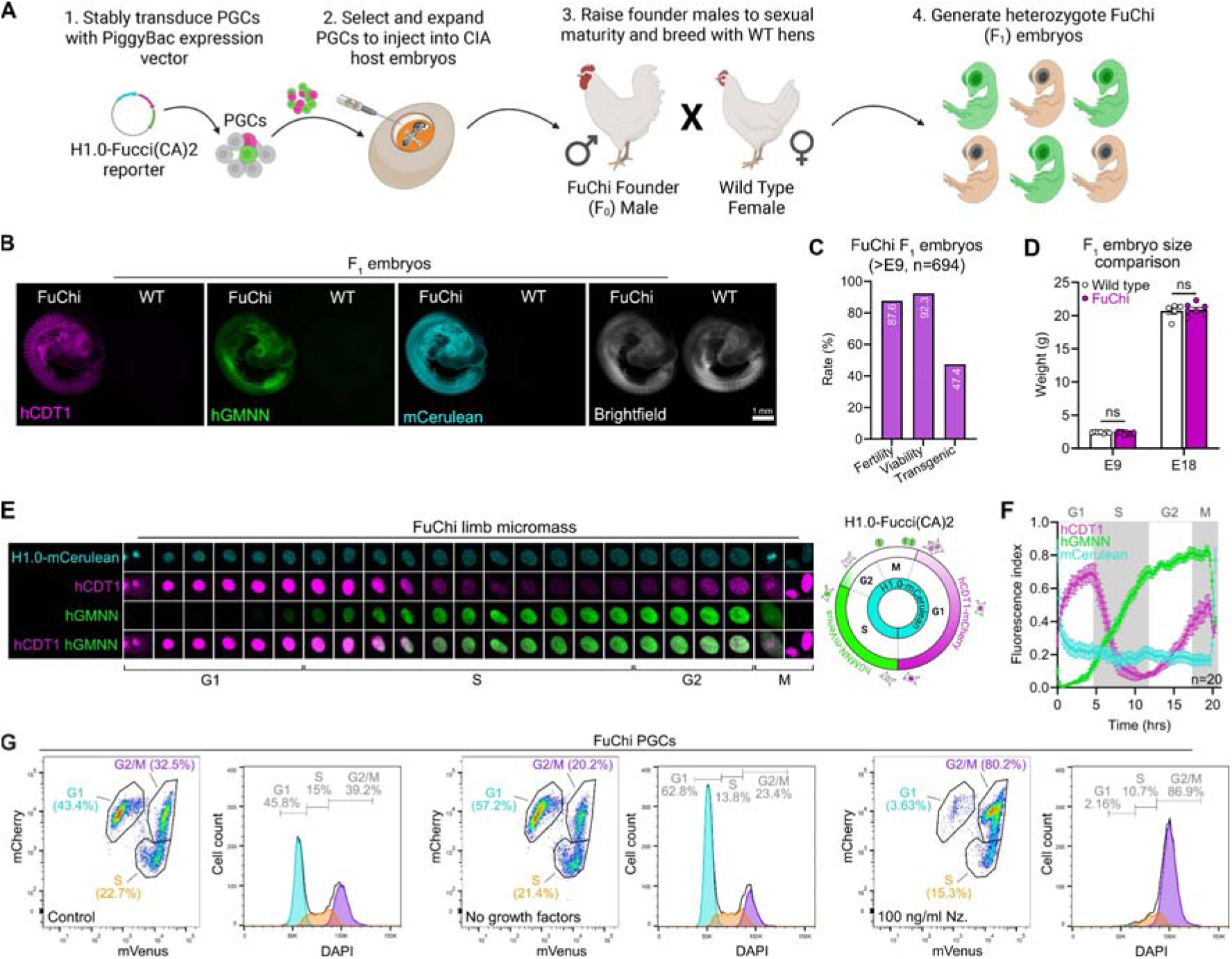
Generation and validation of FuChi reporter chicken. **A)** Schematic of FuChi generation strategy through primordial germ cell (PGC) injection into CIA (chemically induced ablation) sterile host embryos. **B)** H1.0-Fucci(CA)2 expression in FuChi F_1_ embryos compared to wild type. **C)** Fertility, viability and carrier rate of FuChi surrogate hosts. **D)** FuChi embryos develop normally and are comparable in size to wild type embryos. **E)** Individual cell montages from limb micromass cultures derived from FuChi embryos with accompanying schematic of H1.0-Fucci(CA)2 expression profiles. **F)** Temporal normalised fluorescence intensity profile of FuChi limb micromass cells based on live cell imaging data. **G)** Validation of H1.0-Fucci(CA)2 reporter system by flow analysis in PGCs derived from FuChi embryos in both control, growth factor removed, and nocodazole (Nz) treated conditions. The reporter profiles accurately reflect DNA content as determined by DAPI staining in both control and experimental conditions. Scale bar = 1 mm.

To further investigate the operation of the H1.0-Fucci(CA)2 construct in our FuChi chickens we derived PGCs and limb micromass cultures from FuChi F_1_ embryos for flow cytometry analysis and confocal imaging, respectively. The temporal fluorescent profiles of the H1.0-Fucci(CA)2 biosensor in micromass cells closely matched those of our H1.0-Fucci(CA)2-DF1 cells (Figure 2D) with clear G1, S, G2/M populations identifiable through Fucci(CA)2 biosensor abundance with labelling of nuclei by H1.0-Cerulean throughout the cell cycle and a strong H1.0-Cerulean peak evident during M-phase (Figures 3E and 3F; Supplementary video 3). Flow cytometry analysis of three independent PGC lines (two male, one female) confirmed the fidelity of the reporter in both control and experimental culture conditions in which cell cycle progression was perturbed (Figures 3G and S2F). Taken together, these results establish FuChi chickens as the first viable stably expressing avian cell cycle biosensor model for *in vivo* studies.

### Monitoring cell cycle progression throughout chicken embryonic development

Having validated the functionality of H1.0-Fucci(CA)2 in cell lines derived from FuChi embryos we surveyed reporter activity during embryonic development (Figure 4). In Eyal Giyaldi & Kochav (EG & K)^42^ stage X embryos, a large proportion of blastoderm cells were observed in G2/M phase, consistent with them entering diapause during lower storage temperature prior to incubation (Figure 4A)^43^. As gastrulation progresses, the primitive streak and presomitic mesoderm displayed a large proportion of cells in S, and G2/M phase, whereas the developing mesendoderm and neural plate regions contain cells predominantly in G1 (Figure 4B). As development proceeds, FuChi embryos reveal distinct regions of proliferation including the tailbud (Figure 4C) and forming regions of the central nervous system (Figure 4D) which exhibit high levels of proliferation as determined by the GMNN expression profile.

**Figure 4.**
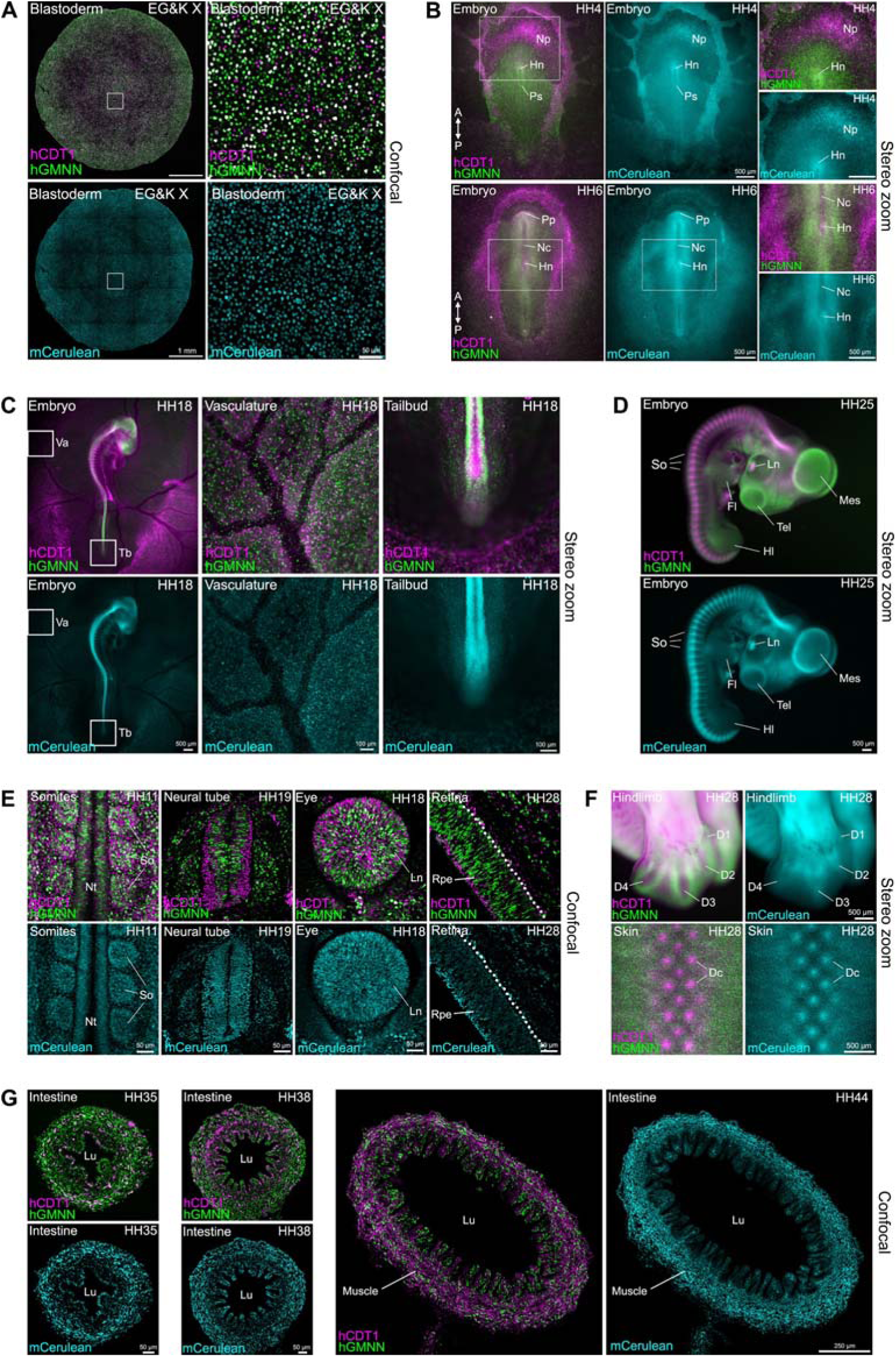
H1.0-Fucci(CA)2 reporter expression in embryonic tissues. H1.0-Fucci(CA)2 biosensor expression in FuChi embryos during early development, visualised using either confocal or fluorescent stereo zoom microscopy. **A)** In the blastoderm there is an abundance of proliferating cells, many of which are double labelled with mCherry-hCDT1/mVenus-hGMNN (S, G2/M). **B)** In early embryos the region of the presumptive neural plate (Np) anterior to Hensen’s node (Hn) and the primitive streak (Ps), is largely non-proliferative (G1; mCherry-hCDT1 positive) as is the notochord (Nc) and precohordal plate (Pp). The anterior (A) – posterior (P) axis is included for orientation purposes. **C)** FuChi embryos also enable visualisation of circulating blood cells during vasculogenesis and the proliferative regions of the tail bud. **D)** As embryonic development continues, distinct proliferation profiles can be seen such as the proliferative developing regions of the central nervous system (mesencephalon (Mes) and telencephalon (Tel)), hindlimb (Hl) and forelimb (Fl), and the non-proliferative somites (So) and lens (Ln). **E)** Embryonic tissues from FuChi embryos visualised using confocal microscopy. Segregated proliferating and non-proliferating populations are present in the somites, neural tube (Nt), lens, and retinal pigmented epithelium (Rpe). **F)** Embryonic tissues of FuChi embryos visualised by stereo microscopy show the stalling of cells into G1 during mesenchymal condensation is visible in structures such as the dermal condensates (Dc) of feather primordia and the developing limb digits (D1-4). **G)** Areas of regional proliferation are established in the developing intestine as villi (Vi) form and line the lumen (Lu). Initially widespread cell division occurs throughout the intestine followed by the outer muscle layers becoming largely non-proliferative and cell division mainly occurs in the epithelium. Scale bars as annotated.

Static and live imaging of developing somites revealed that their epithelial cells undergo extensive proliferation, with a high proportion of S, and G2/M cells, compared to the mesenchyme, which is predominantly in G1 as they undergo condensation (Figures 4D and 4E; Supplementary video 4). Cryosections of fixed FuChi retinal tissue, a tissue known to be highly proliferative during development^44^, displayed a high proportion of cells in S, G2/M (Figure 4E). Similarly, other sites of mesenchymal condensations such as the developing digits and feather buds show an abundance of G1-labeled cells (Figure 4F). Analysis of the developing intestine across several developmental stages revealed proliferative changes as the muscle layers developed, whilst also confirming the presence of proliferative epithelial cells in the base of the crypts at later stages consistent with a stem cell, and subsequent transit amplifying cell population, arising during late embryonic development^45,46^ (Figure 4G).

Taken together, these results confirm that the intensity profiles of the H1.0-Fucci(CA)2 biosensor in FuChi embryos illuminate the known cell cycle profiles of those tissues.

### Robust H1.0-Fucci(CA)2 stability in fixed tissues and immunodetection after paraffin embedding

Next, to assess the stability and usability of H1.0-Fucci(CA)2 in FuChi tissues we analysed persistence of the fluorescence signal following common methods of fixation and tissue preparation (Figure S3). H1.0-Fucci(CA)2 signal was stable following a range of fixation methods, compared to unfixed specimens (Figure S3A). We observed only a slight decrease in the signal to noise ratio due to increased autofluorescence of the fixed samples (Figure S3B).

While H1.0-Fucci(CA)2 signal can be directly observed in cryosections and fixed wholemount tissue (Figure 4), we wanted to test whether our reporter embryos could be combined with gene and protein expression analysis. We performed fluorescent *in situ* hybridisation to detect expression of the hedgehog target gene *PTCH1* in the developing embryo (Figure 5A). *PTCH1* is expressed in neural tube, limb bud, and somites, with expression closely correlating with proliferating cells in the neural tube (Figure 5A)^47^. Conversely, *SHH* expression, which induces *PTCH1* expression, is restricted to the notochord and floor plate of the neural tube (Figure 5B)^48^. Staining of FuChi embryonic spinal cord cryosections with a YAP1 antibody (Figure 5C) revealed that expression is localised in the progenitor zone, a region associated with high proliferation and regulated by YAP^49^. Additionally, the inclusion of epitope tags in H1.0-Fucci(CA)2 also enables detection of antigens by immunofluorescence in formalin fixed paraffin embedded (FFPE) sections and cryosections of embryonic kidney (Figure 5D).

**Figure 5.**
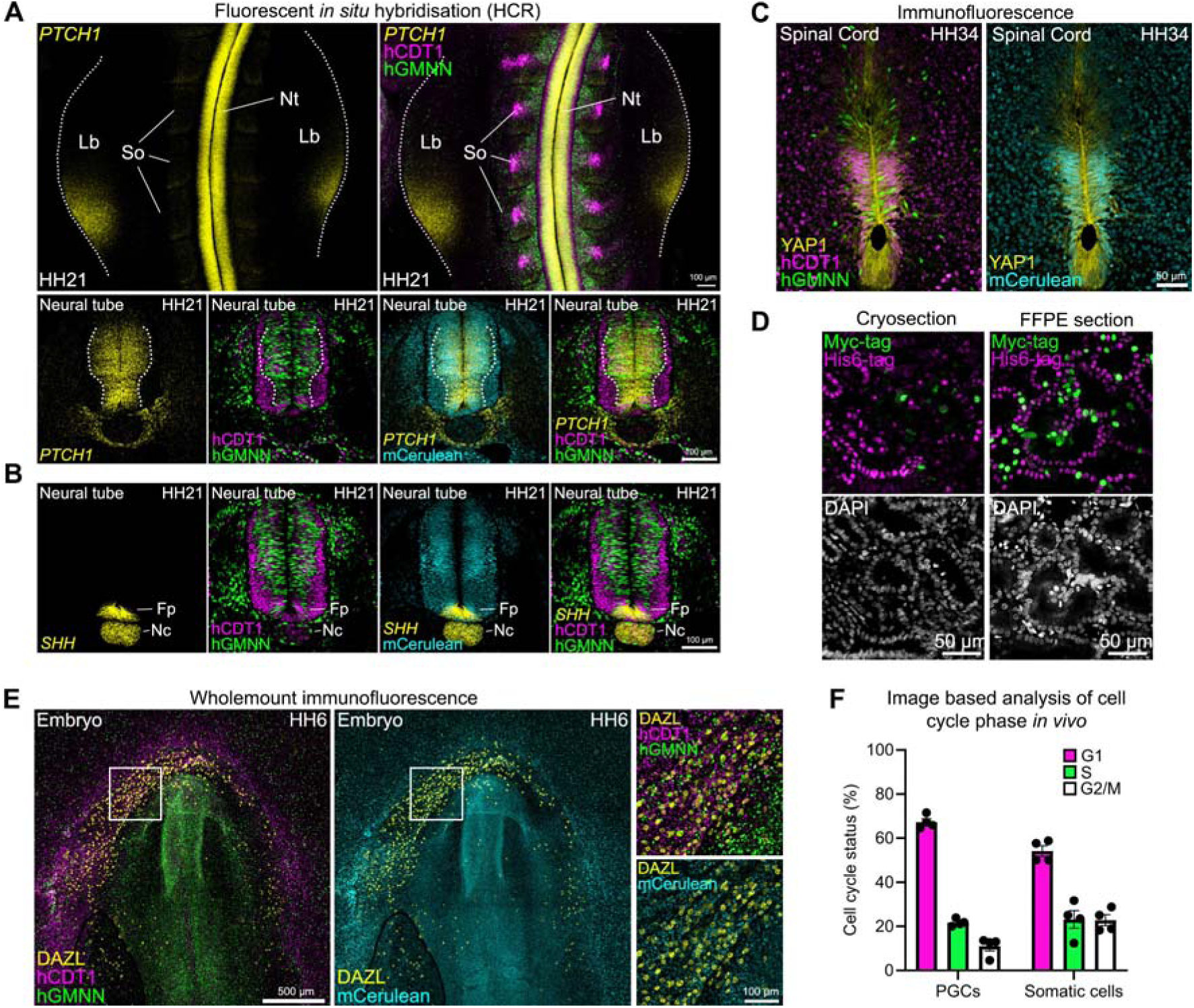
Combining cell cycle status with detection of endogenous protein and mRNA expression. **A)** Hybridisation Chain Reaction (HCR) to detect *PTCH1* transcript expression in HH18 FuChi embryos. Top panels show wholemount staining and bottom panels are cryosections of these embryos. Strong *PTCH1* mRNA expression can be seen in the neural tube (Nt; demarcated by dashed lines), corresponding with a region of high proliferation, in the limb bud (Lb), and with weak expression in the somites (So). **B)** Expression of *SHH*, which induces *PTCH1* expression, is restricted to the floor plate (Fp) and the notochord (Nc). Expression of the fluorescent reporters remains robust following HCR. **C)** Immunofluorescent detection of total YAP1 protein in a cryosection of the developing spinal cord of a HH34 FuChi embryos. **D)** Examples of cryo-embedded and formalin fixed paraffin embedded (FFPE) sections of HH29 mesonephros with immunofluorescent detection for the epitope tags contained in the FuChi construct (His6-mCherry-hCDT1, Myc-mVenus-hGMNN). Fluorescent profiles are comparable between the two section preparation types. **E)** Expression of FuChi reporter in migrating PGCs, identified by wholemount immunofluorescence for the germ cell marker DAZL, as they move anteriorly towards the germinal crescent. **F)** Percentage of PGCs and neighbouring somatic cells in each cell cycle phase determined from confocal images (Between 55 and 172 of each cell type were analysed per embryo (n = 4 embryos). Scale bars as annotated.

To further explore the utility of FuChi embryos, we tested whether it was possible to assess the proliferation profile of native migrating cells using immunofluorescence on wholemount embryos (Figures 5E). In chickens, PGCs migrate anteriorly from the centre of the blastoderm, ahead of the forming embryo to populate the germinal crescent during early development. Here they enter the bloodstream and are translocated to the anterior vitelline veins where they exit the blood stream and migrate into the genital ridges, before finally populating the gonads^50,51^. However, the proliferative status of PGCs during their initial migratory steps, prior to entering the bloodstream, is unknown. After establishing parameters (see methods for more details; Figure S4) for our imaging-based approach, we found that as PGCs migrate towards the germinal crescent the majority of cells reside in G1 (67.3%), compared to those in S (21.7%) and G2/M (11.0%) (Figure 5F). This differs from both neighbouring somatic cells (Figure 5F) and also from PGCs in culture (Figures 3G, S2F, and S4), indicating that unlike the rapidly proliferating germ cells in the gonads, PGCs at these initial migratory stages are not undergoing frequent cell division.

### Real time visualisation of cell cycle dynamics during gastrulation

A major attraction of Fucci systems is that they can be used to image cell cycle progression, not only in isolated live cells, but also in complex tissues and even whole embryos. Control of the cell cycle plays an important role in the embryonic development of all organisms, including chick. However, so far there is only limited information available on cell cycle changes during avian development. To assess the potential of the H1.0-Fucci(CA)2 system for live imaging we used lightsheet microscopy to monitor cell cycle changes from the onset of development after egg laying (EGK XII) to the primitive streak stage (HH4) (Figure 6A, Supplementary video 5). Embryos were imaged at 10x magnification, allowing visualisation and detection of mVenus (hGMNN) and mCherry (hCDT1) in all nuclei of the surface layers of the embryo at 8-10 minute time intervals. To quantitatively assess changes in the distribution of cells in different cell cycle phases during the early stages of development we measured the total number of nuclei detected in G1, S, and G2/M based on hGMNN-mVenus and hCDT1-mVenus fluorescence as a function of time in both the embryonic and extra embryonic territories (Figure 6B). This revealed that at the onset of development most cells are in S-phase and progress into G2 followed by mitosis and G1 in a partially synchronised manner which persists in a further subset of cells during primitive streak formation (Figure 6B). Next, to characterise the temporal cell cycle dynamics *in vivo* we measured the kinetics of fluorescence changes in individual cells in the embryonic and extraembryonic regions at the onset of development (Figure 6C, Supplementary videos 6-8). This analysis revealed that the cell cycle is around 10 hrs in both territories (Figure 6D; representative of three independent repeats). The synchronised manner of cells progressing from S through G2 followed by mitosis to G1 from the onset of development was also apparent in the global analysis of cells in the specific regions analysed (Figure S5A).

**Figure 6.**
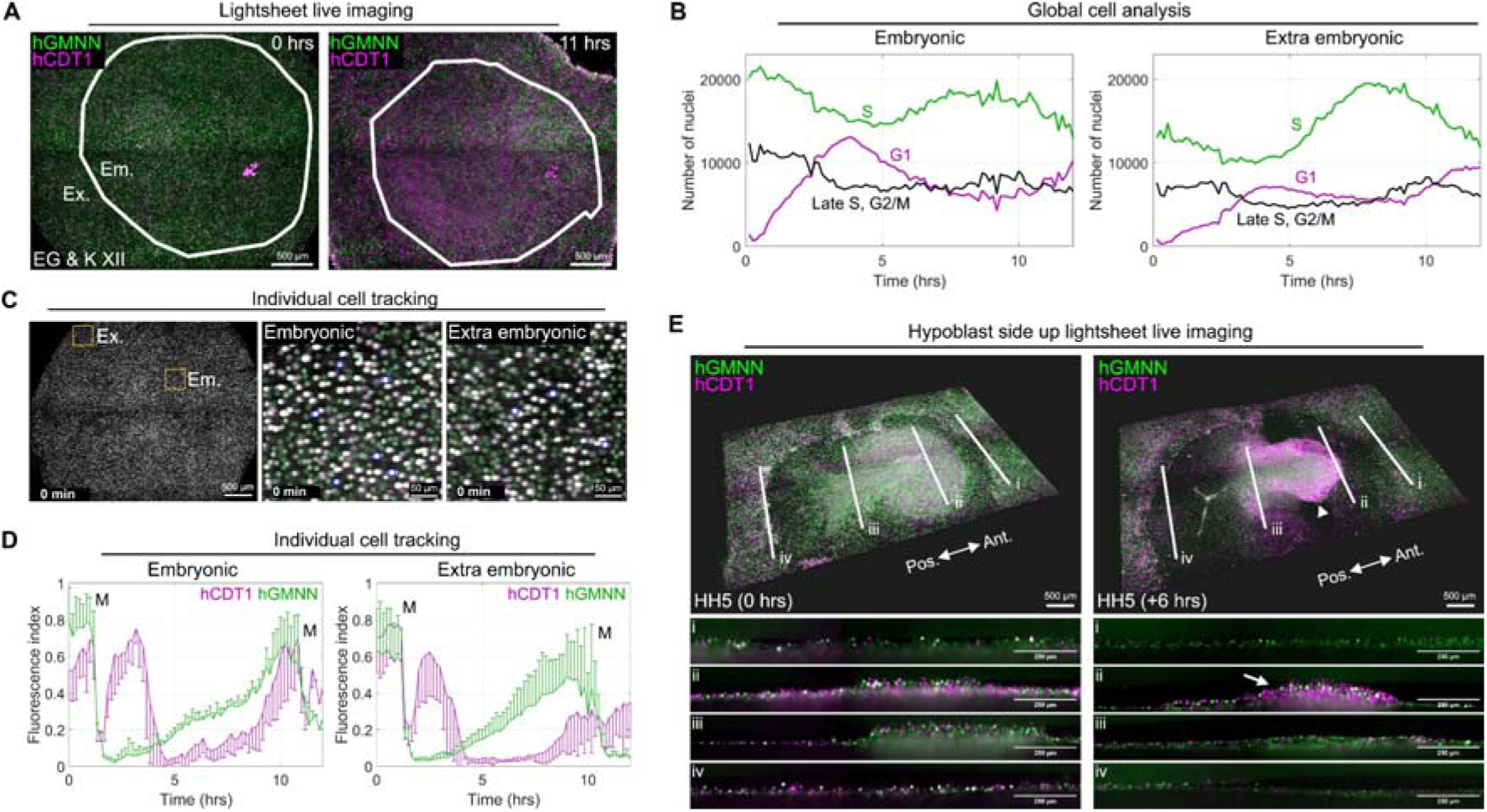
Visualisation and assessment of cell cycle dynamics during early embryonic development. **A)** Epiblast surface views from a representative lighsheet experiment performed on a FuChi embryo at the start and after 11 hours of development. The boundary separating the embryonic (Em.) and extra embryonic regions (Ex.) is indicated by the white line, calculated as described in methods. **B)** Global cell counts of embryonic and extra embryonic regions of the FuChi embryo visualised by lightsheet microscopy across time in Figure 5A. All cells detected in the analysed regions were categorised based on their fluorescence profile at the time interval; hCDT1-mCherry positive (G1; magenta), hGMNN-mVenus positive (S; green), or positive in both channels (Late S, G2/M; black). The same observations were made across four independent lightsheet experiments. **C)** Left panel: Representative image of epiblast surface view from light sheet timelapse imaging of a FuChi embryo with regions of interest in the embryonic and extra embryonic regions highlighted (yellow boxes). Right panels: Individual cells (blue rings) were tracked in the embryonic and extraembryonic regions. Images represent a single time point from a lightsheet timelapse video. **D)** Normalised fluorescence profiles of individual cells in embryonic and extra-embryonic cut out regions highlighted in (C). Error bars represent standard deviation. **E)** 3D hypoblast view of a stage HH5 FuChi embryo at two successive time points 6 hrs apart. Four optical sections (panels below) taken along the anterior (Ant.) to posterior (Pos.) plane, including both the extraembryonic (i) and embryonic regions (ii-iv) are also shown at positions indicated by white lines. Note that cells egressing from the streak (arrow) and cells of the prechordal plate and anterior mesoderm cells already migrated away (arrowhead) are predominantly in the G1. Scale bars as annotated.

Finally, the epiblast view showed that during primitive streak formation many cells in the deeper layer of the embryo resided in G1 while migrating away from the streak (Figure S5B). To investigate this in more detail we performed hypoblast side up experiments (Figure 6E, Supplementary video 9), which allows better viewing of hypoblast cells and cells emerging from the streak after their ingression. Strikingly, these experiments (three independent repeats) revealed that mesendoderm cells emerging from the primitive streak are in S phase, transition through G2/M, and enter G1 during their migration away; this is particularly evident in cells forming the prechordal plate (Figure 6E, Supplementary video 9). These whole embryo live imaging results demonstrate that the H1.0-Fucci(CA)2 system allows tracking of cell cycle progression *in vivo* for prolonged periods of time. Importantly, it highlights the dynamic cell cycle changes that cells undergo during key morphological events such as gastrulation. The significance of these findings, including to the extent to which cell cycle progression influences cell fate, will be elaborated in more detail in future experiments on FuChi embryos.

## Discussion

Here we describe H1.0-Fucci(CA)2 a novel tricistronic cell cycle biosensor capable of resolving G1, S, G2 and M cell cycle phases from a single construct. Further we describe the first avian cell cycle reporter model and demonstrate the functionality of the H1.0-Fucci(CA)2 biosensor in chickens *in vivo*. Fucci(CA)2 functions corectly in the chicken both *in vivo* and *in vitro*, displaying a fluorescent reporter profile consistent with that observed in mammalian systems^9^. A major advantage of this reporter is its ability to distinguish between S and G2/M phases - a key improvement over the original Fucci(SA)2 system. In addition, the inclusion of H1.0-mCerulean allows for continuous cell tracking throughout the cell cycle and enables specific identification of M-phase via labelling of mitotic chromatin. The incorporation of epitope tags further expands the system’s utility by facilitating analysis in fixed tissues.

Although Fucci(SA)2 systems using human CDT1 fusion proteins have previously been used in chicken cells *in vitro* and through transplantation *in vivo*^22,23^, our results suggest that in this system human CDT1 is not efficiently degraded during S phase, thereby reducing the overall fidelity of the Fucci(SA)2 reporter in chick. This likely arises from differences between the core residues of the Cy motif in mammalian (Arg, Arg, Leu) and avian (Lys, Arg, Leu) CDT1, which are essential for SCF-mediated CDT1 degradation^3,52,53^. This is consistent with human CDT1 based-Fucci reporters requiring SCF degradation not being functional in zebrafish^38^. The Fucci(CA)2 system overcomes species differences in the Cy motif by utilising the PIP box of CDT1, a feature conserved across metazoans^54^. As hGMNN degradation is well conserved and functional between species^38^, our data suggest that the pre-existing human Fucci(CA)2 reporter system will function in other model organisms, without the need for the generation of new species-specific CDT1 variants.

The ability to analyse cell cycle status *in vivo* in both real time and in fixed sections enables many downstream applications. Here, we investigated the proliferative status of migrating PGCs during early embryonic development *in vivo*. Previous studies analysed chicken PGC cell cycle status either in culture^23,55^ or from stages beyond E2.5^56,57^, at which point the PGCs have entered the bloodstream. Our results reveal that at HH stage 6, as PGCs migrate anteriorly to the germinal crescent and prior to entering the bloodstream, the majority of cells are in G1 (∼70%). A potential reduction in cell division before entering the vasculature is consistent with PGC number not rapidly increasing across these stages^58^, as well as PGC cultures derived from EG & K stage X embryos showing reduced proliferation compared to those derived from circulating PGCs^55^. This suggests a trade-off between proliferation and active migration in PGCs at this stage, distinct from earlier passive embryo-wide movements or later circulation via the bloodstream^59^. Beyond germ cell development, FuChi embryos could be used to interrogate the proliferative status of other migrating cells such as neural crest cells, leveraging the chick embryo’s well-established advantages for neural crest research^60,61^.

The cell cycle duration observed in the embryonic and extra embryonic tissues here is faster than that measured in tissue culture and is consistent with previous reports from tracking experiments in gastrulating embryos^29^. The results show that most cells are arrested in S phase during the diapause that follows egg laying. Upon re-entering development, cells complete S-phase and synchronously enter G2, followed by mitosis, resulting in the observed partial synchrony of later cell divisions. The results also suggest the possibility that cells are in S phase while emerging from the primitive streak and transition to G1 as they migrate away from the streak, which is particularly prominent for cells forming the prechordal plate and lateral plate mesoderm. The significance of this pattern and its link to differentiation could be elucidated by detailed cell tracking at higher magnification in future work.

Developing Fucci-expressing cell lines requires strategies that achieve stable integration of the construct at moderate expression levels with balanced stoichiometry of the fluorescent reporters. Previously, this was achieved using the Flp-In system^13,62^. In this study, we successfully applied PiggyBac transposon technology in the chick, consistent with recent results in human cells^24^. This approach achieves comparable integration without the need for prior genomic insertion of a Flp-In cassette or reliance on viral transduction. As such, the H1.0-Fucci(CA)2 PiggyBac construct described here should enable straightforward generation of Fucci reporter lines in diverse systems including primary cells, organoids, and cancer cell models across mammalian and avian species, as well as alternative animal models such as mouse and zebrafish.

The chick embryo is a powerful model system that has been instrumental in advancing our understanding of vertebrate development and disease since the 17th century^63^. Over the past two decades, progress in chicken stem cell and transgenic technologies^26,27,64,65^ has enabled the generation of several fluorescent reporter and gene knockout chicken lines, leading to important discoveries in embryonic development, human congenital disorders, and infectious diseases^32,66–70^. The growing repertoire of transgenic tools - now including the FuChi line - is helping to establish the chicken as a highly tractable alternative to more commonly used models, such as the mouse and zebrafish. Furthermore, the chorioallantoic membrane (CAM) of FuChi embryos will provide a platform to investigate host tissue proliferation and dedifferentiation in response to human or animal xenografts (tumours or cell lines) or viral infection, and to evaluate potential therapies in a live, vascularized microenvironment^71^.

From a 3Rs (Replacement, Reduction and Refinement) perspective, the chick embryo offers clear ethical and logistical advantages. For example, in both the United Kingdom and European Union, embryos up to embryonic day 14 are not classified as protected animals, and their use does not require the sacrifice of a pregnant female. This supports Reduction in the number of protected animals, as a single small transgenic flock can generate ∼1,300 embryos annually^72^. The chick also contributes to Replacement of early-stage mammalian models and allows for Refinement through non-invasive *in ovo* imaging and precise manipulation during development.

## Limitations

While the FuChi system offers comprehensive resolution of all major cell cycle phases (G1, S, G2, and M) and enables high-resolution live imaging of proliferation during early avian development, several limitations remain. First, although FuChi expression is strong and widespread in early embryos, its performance in later developmental stages has not yet been systematically evaluated. Differences in chromatin environment, epigenetic silencing, or protein turnover across tissues and timepoints could affect biosensor performance at later stages. Second, although FuChi embryos develop normally, some adults showed deleterious effects, likely due to piggyBac-mediated insertion-site variability, highlighting the need for a chicken safe-harbour locus to enable ubiquitous expression while minimising insertional risk. Third, as previously reported, the Fucci(CA) probes rely on the CUL4^DBB1^mediated degradation pathway; therefore, the intensity of the hCDT1(1/100)Cy(-) signal may be reduced in contexts where DNA damage is induced (e.g., by UV irradiation). Fourth, although Fucci(CA)2 distinguishes G1 from S, and S from G2/M, and the mCerulean-tagged histone H1.0 enables visualisation of mitotic chromatin, precise quantification of transitions (e.g., G2 to M) may still require complementary temporal data or additional markers, particularly in highly dynamic contexts. Finally, although the FuChi biosensor enables clear phase discrimination, analysing proliferation in rapidly morphing or migrating tissues (e.g., the primitive streak or neural crest streams) may require high-frame-rate imaging or integration with lineage markers to avoid misinterpreting positional shifts as cell-cycle transitions. This will be enabled by application of the latest Ai-driven segmentation and tracking algorithms^73,74^.

## Conclusion

In conclusion, the FuChi transgenic line provides a powerful and versatile tool for real-time analysis of the cell cycle in the chicken embryo, both *in vivo* and *in vitro*, across multiple developmental stages. We show that the Fucci(CA)2 system functions robustly in the avian context, offering precise discrimination of all major cell cycle phases—including mitosis—while the addition of H1.0-mCerulean and epitope tags enhances both live imaging and fixed-tissue analysis, including in FFPE samples. The stability and accessibility of this reporter line make it a valuable resource for researchers worldwide. Collectively, these advances position FuChi embryos as a premier *in vivo* system for investigating developmental cell-cycle kinetics, with clear technological advantages over current mouse models. We anticipate that FuChi will advance studies in development, growth, tissue homeostasis, and disease. FuChi eggs are now available upon request from the National Avian Research Facility at the Roslin Institute, UK.

## Acknowledgements

We would like to thank all of the staff belonging to the NARF and Bioimaging units of the Roslin Institute without whom this project would not have been possible. This research was funded by BBSRC (BBS/E/D/10002071; BBS/E/RL/230001C) and NC3Rs (NC/T002328/1).

Edinburgh Genomics are supported with core funding from the Natural Environment Research Council (UKSBS PR18037). Figure schematics were created using BioRender.

## Declaration of generative AI and AI-assisted technologies in the manuscript preparation process

During the preparation of this work the author(s) used ChatGPT in order to assist in grammar checks, and rephrasing of paragraphs for conciseness. After using this tool/service, the author(s) reviewed and edited the content as needed and take(s) full responsibility for the content of the published article.

## Supplementary Figures

**Supplementary Figure 1.**
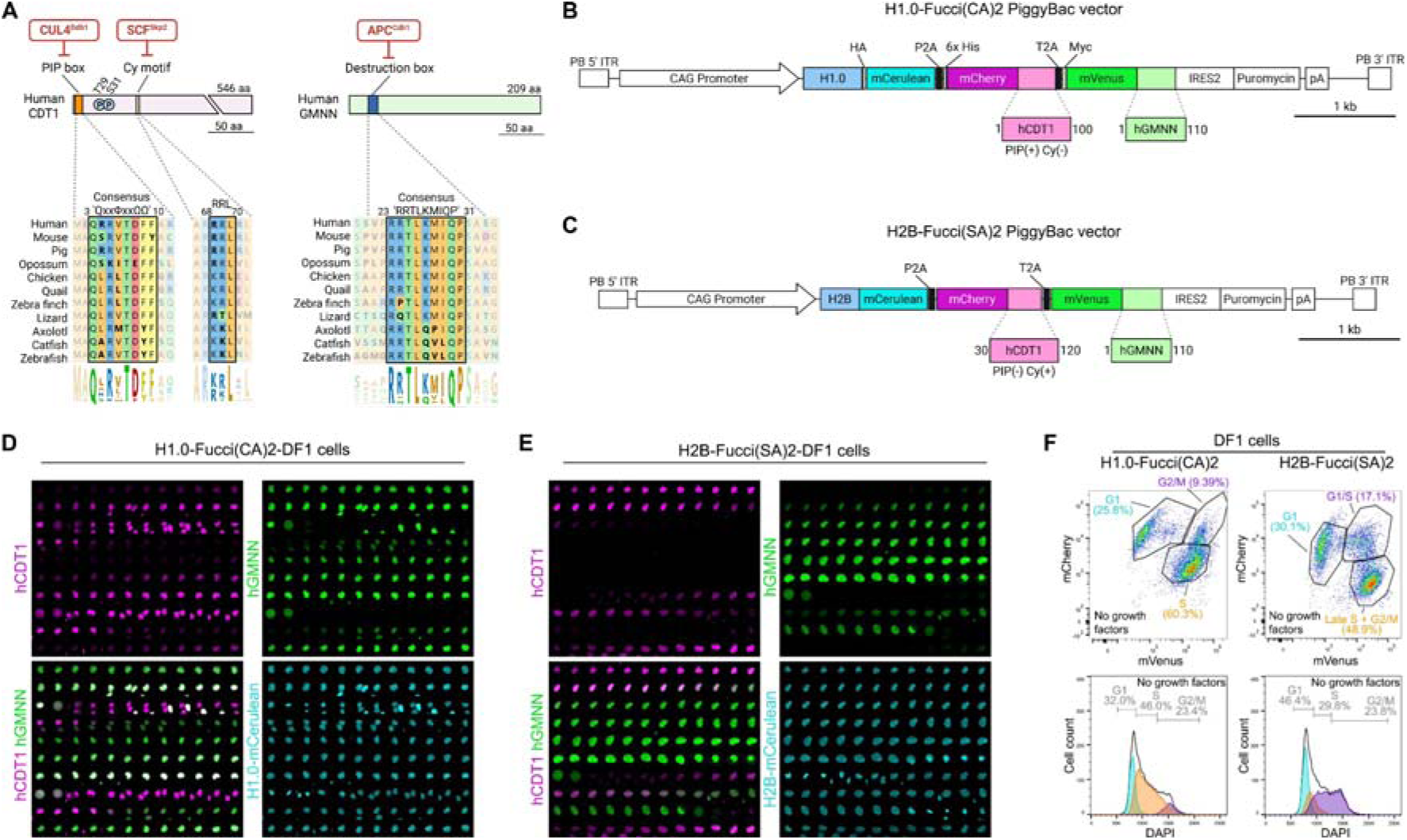
Validation of Fucci reporter systems in chicken cells *in vitro* and *in vivo*. **A)** Comparison of the PIP box (targeted by CUL4) and Cy motif (targeted by SCF) degradation domains of CDT1 and the destruction box (targeted by APC) of GMNN across vertebrate species. While the destruction box of GMNN is well conserved in mouse, human and chicken, as are the core residues of the PIP box (Ψ represents any moderately hydrophobic amino acid (L, V, I, or M); Ω is an aromatic residue (Y or F)), the Cy motif of CDT1 is not well conserved across species. **B and C)** Design of PiggyBac Fucci reporter constructs. **B)** The H1.0-Fucci(CA)2 reporter cassette uses a CAG promoter to drive expression of H1.0-mCerulean, hCDT1(1/100)PIP(+)Cy(-)-mCherry, and hGMNN(1/110)-mVenus fusion proteins, each containing a 5’ epitope tag (HA, 6x His, Myc), and which are upstream of puromycin resistance gene. **C)** The H2B-Fucci(SA)2 reporter cassette expresses H2B-mCerulean, hCDT1(30/120)PIP(-)Cy(+)-mCherry and hGMNN(1/110)-mVenus fusion proteins, upstream of puromycin. ITR = Inverted terminal repeats. The presence or absence of the PIP domain and Cy motif of human CDT1 proteins are denoted in parentheses. **D)** Extended timelapse images showing two full cell cycles of H1.0-Fucci(CA)2 expressing DF1 cells. **E)** Extended timelapse images showing two full cell cycles of H2B-Fucci(SA)2 expressing DF1 cells. **F)** Comparison of Fucci(CA)2 vs Fucci(SA)2 reporter systems in chicken DF1 fibroblasts. In the absence of serum, an increased proportion of cells are observed in G1 and S phase through DNA content analysis using flow cytometry. This shift is accurately captured in the fluorescent profiles of H1.0-Fucci(CA)2 but not H2B-Fucci(SA)2 cells. For comparison, control conditions are shown in Figures 1F and 1I. **C)** Table showing fertility, viability and transgenic rate of FuChi embryos across various developmental stages. **D)** FuChi chicks can be hatched, are viable, and reporter expression can be detected. **E)** Validation of H1.0-Fucci(CA)2 reporter system by flow analysis in two independently derived PGC lines derived from FuChi embryos in both control, growth factor removed, and nocodazole (Nz) treated conditions. The reporter profiles accurately reflect DNA content as determined by DAPI staining in both control and experimental conditions. Results from a third PGC line (#2) can be seen in Figure 2G.

**Supplementary Figure 2.**
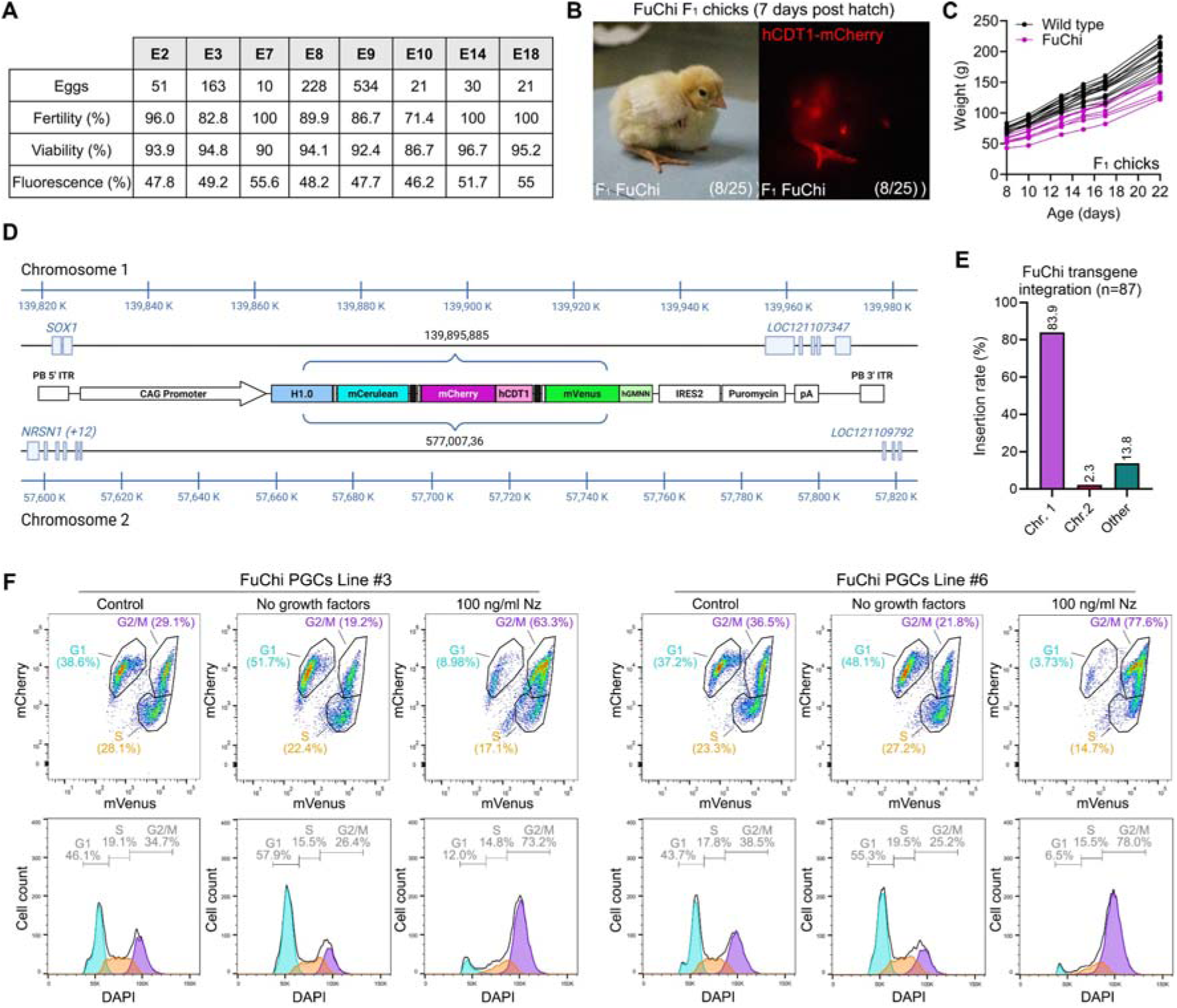
Further characterisation of the viability and functionality of FuChi chickens. **A)** Table showing fertility, viability and transgenic rate of FuChi embryos across various developmental stages. **B)** FuChi chicks can be hatched, are viable, and reporter expression can be detected. **C)** Hatched FuChi chicks are smaller than wild type chicks from the same bath but gain weight at a similar rate. **D)** Mapped transgene integration sites in FuChi chickens based on analysis of a pool of five embryos. **E)** Percentage of chromosome integration sites identified by sequencing present in FuChi embryos. At least one more transgene insertion is present that was not present in the embryos analysed in (D). In total, 87 embryos were analysed at two developmental stages (HH19; n = 32; HH35; n = 55). **F)** Validation of H1.0-Fucci(CA)2 reporter system by flow analysis in two independently derived PGC lines derived from FuChi embryos in both control, growth factor removed, and nocodazole (Nz) treated conditions. The reporter profiles accurately reflect DNA content as determined by DAPI staining in both control and experimental conditions. Results from a third PGC line (#2) can be seen in Figure 3G.

**Supplementary Figure 3.**
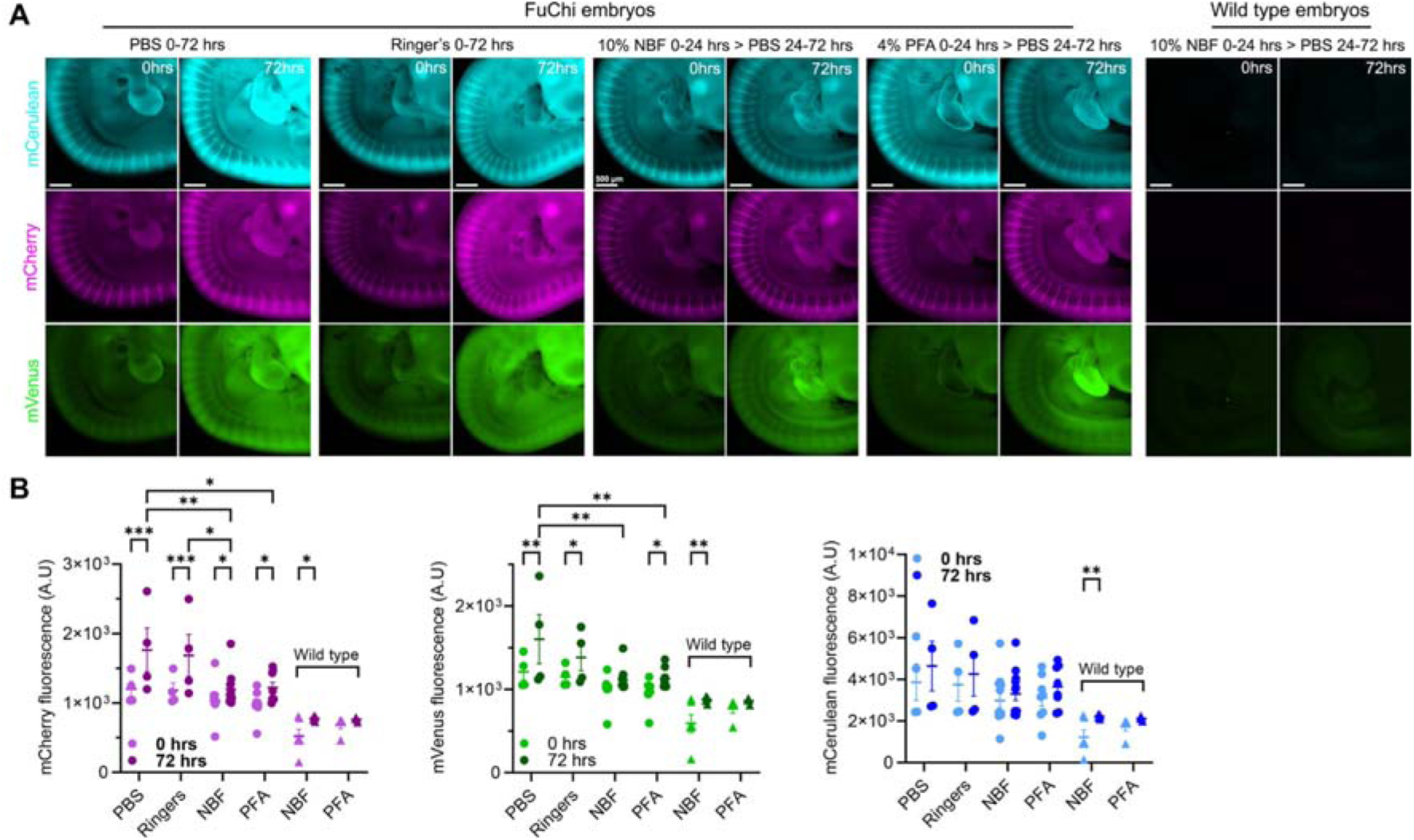
Reporter fluorescence quality post-fixation. **A)** Example images of FuChi or wildtype embryos which were fixed and/or stored in each medium condition as annotated and imaged at 0 and 72 hours. **B)** Fluorescent reporter levels at 0 and 72 hrs in FuChi and wild type embryos following fixation and/or storage. Fixation in 10% neutral buffered formalin (NBF) or 4 % paraformaldehyde (PFA) stabilises H1.0-Fucci(CA)2 reporter expression while reducing increase in autofluorescence levels observed without fixation. Error bars represent S.E.M. *P<0.05, **P<0.01, ***P<0.001; Two-Way ANOVA with pairwise comparisons. Scale bars as annotated.

**Supplementary Figure 4.**
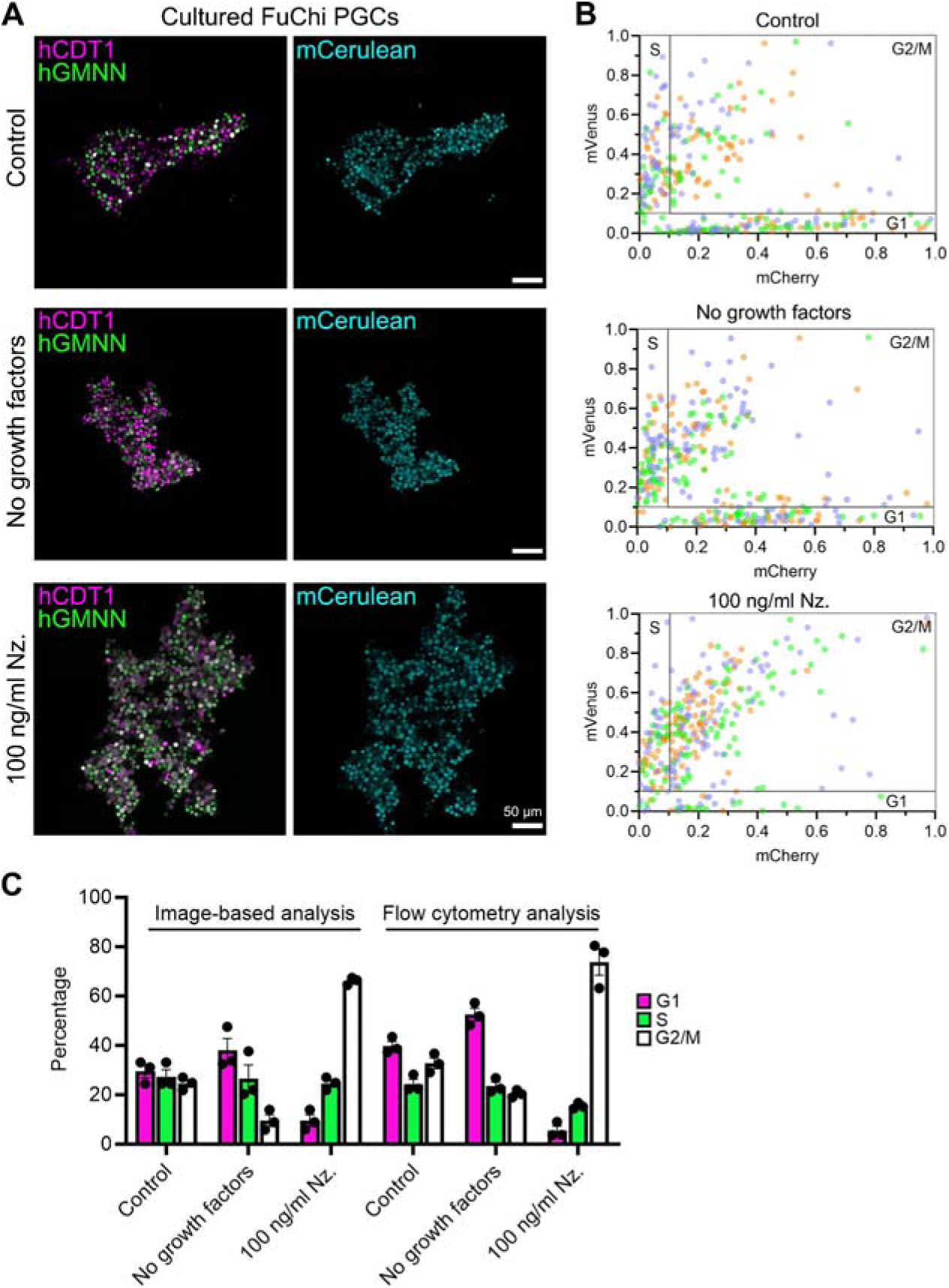
Parameterisation for image-based analysis of H1.0-Fucci(CA)2 reporter in FuChi cells *in vitro* and *in vivo*. **A)** Representative confocal images of PGCs cultured under different conditions used for analysis of cell cycle stage. The same experimental cells that were analysed by flow cytometry (Figures 2G and S1D) were used. **B)** Scatter plot of relative mCherry and mVenus values from image analysis. Gates for each cell cycle are marked. Orange, green and blue dots represent cells from three independently derived FuChi PGC lines. **C)** Quantification of cell cycle state in PGCs by image based and flow cytometry methodologies. Three independently derived FuChi PGC lines were analysed, with 100 PGCs analysed for image-based quantification. Nz. = Nocodazole. Scale bars as annotated.

**Supplementary Figure 5.**
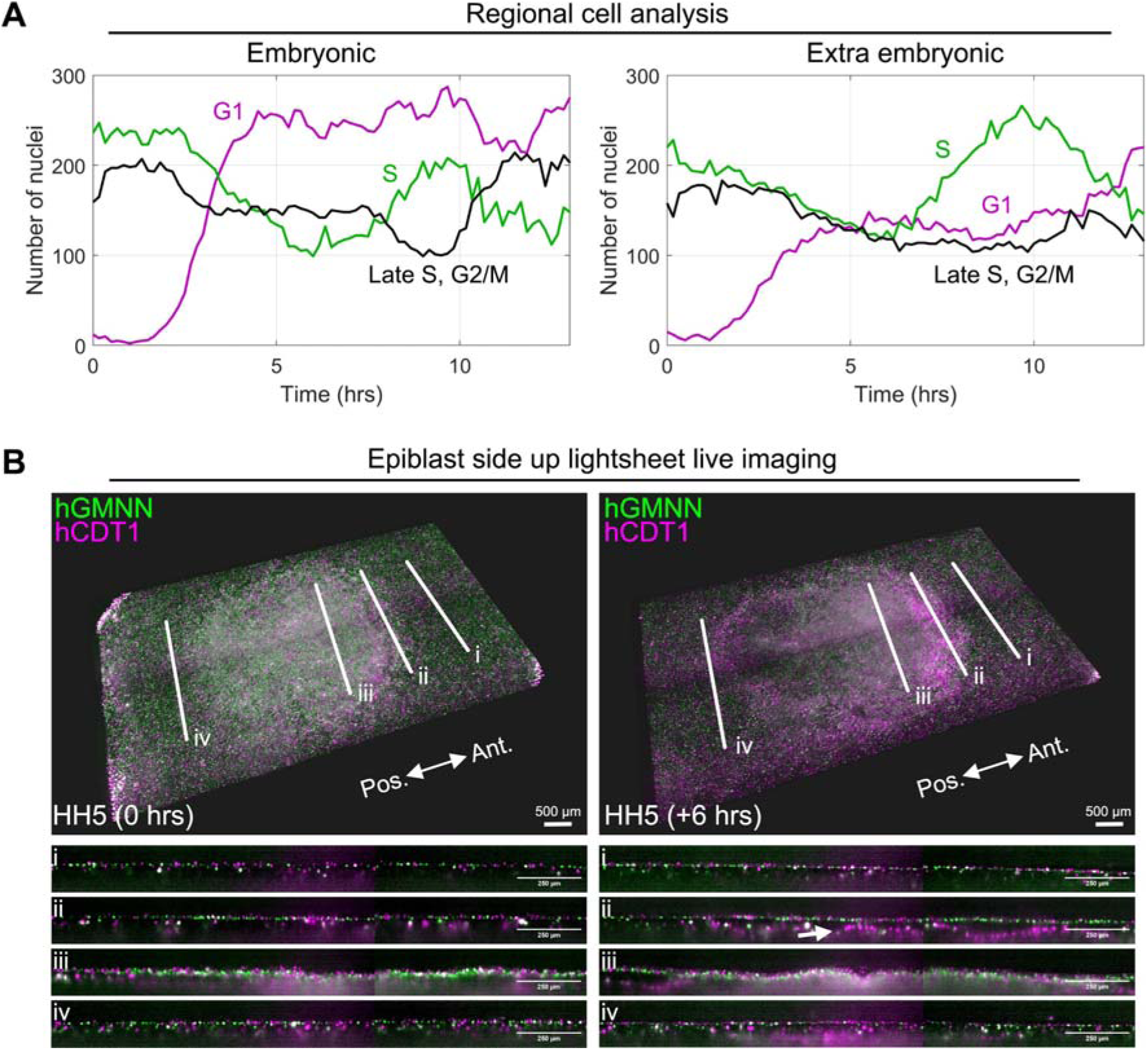
Cell cycle analysis in developing FuChi embryos using lighsheet microscopy. **A)** Cell counts of embryonic and extra embryonic selected regions of the FuChi embryo visualised by lightsheet microscopy across time in Figure 5C. All cells detected in the analysed regions were categorised based on their fluorescence profile at the time interval; hCDT1-mCherry positive (G1; magenta), hGMNN-mVenus positive (S; green), or positive in both channels (Late S, G2/M; black). These results are representative of three independent repeats. **B)** 3D epiblast view of a HH stage 5 FuChi embryo at two successive time points 6 hrs apart. Four optical sections (i-iv) taken from anterior (Ant.) to posterior (Pos.) are also shown at positions indicated by white lines. Note the difference in cell cycle status (G1) of cells egressing from the streak (arrow). Scale bars as annotated.

## Supplementary video legends

**Supplementary video 1**: Imaging of DF1 cells expressing the H1.0-Fucci(CA)2 biosensor. These cells express H1.0-mCerulean (cyan), hCDT1(1/100)Cy(-)-mCherry (magenta), and hGMNN(1/110)-mVenus (green) fusion proteins. Images were captured every 10 minutes by confocal microscopy. Scale bar = 50 µm.

**Supplementary video 2:** Imaging of DF1 cells expressing the H2B-Fucci(SA)2 biosensor. These cells express H2B-mCerulean (cyan), hCDT1(30/120)Cy(+)-mCherry (magenta), and hGMNN(1/110)-mVenus (green) fusion proteins. Images were captured every 10 minutes by confocal microscopy. Scale bar = 50 µm

**Supplementary video 3:** Visualisation of H1.0-Fucci(CA)2 biosensor expression in limb micromass cell cultures derived from a FuChi embryo. hCDT1-mCherry (magenta), hGMNN-mVenus (green), and H1.0-mCerulean (cyan) show dynamic expression through cell division. Images were captured every 10 minutes by confocal microscopy. Scale bar = 50 µm

**Supplementary video 4:** Visualisation of somite formation in a FuChi embryo. Left panel: hCDT1-mCherry (magenta) and hGMNN-mVenus (green) expression. Right panel: H1.0-mCerulean (cyan) reporter expression. Images were captured every 10 minutes by confocal microscopy. Scale bar = 100 µm

**Supplementary video 5:** Lightsheet imaging of a developing FuChi embryo. Changes in cell cycle dynamics in the epiblast during development from stage EGK XIII to HH4. Time interval between images is -8 min and the scale bar represents 500 µm.

**Supplementary video 6:** The dynamic position of selected 330x330 µm regions in the embryonic and extra embryonic areas selected for analysis. The regions (yellow squares) were tracked over time using the velocity field calculated using particle image velocimetry. The time interval between images is 10 mins. Scale bar = 500 µm.

**Supplementary video 7:** Cutout of small epiblast region from the embryonic area of a developing FuChi embryo. Coloured circles indicate detection of hGMNN-mVenus (green) or hCDT1-mCherry (magenta) fluorescence. Manually tracked cells are outlined by blue squares. Scale bar = 50 µm.

**Supplementary video 8:** Cutout of small epiblast region from the extra embryonic tissue of a developing FuChi developing embryo. Coloured circles indicate detection of hGMNN-mVenus (green) or hCDT1-mCherry (magenta) fluorescence. Manually tracked cells are outlined by blue squares. Scale bar = 50 µm.

**Supplementary video 9:** Hypoblast side view of the hypoblast and emerging mesendoderm in a FuChi embryo developing from stage HH3 to stage HH5 with a depth of view of 60 µm. Time interval between images is 10 minutes. Scale bar = 500µm

