## Supplementary figures and images for "FuChi: A cell cycle biosensor for investigating cell-cycle kinetics during avian development"

### Supplementary Figure 1

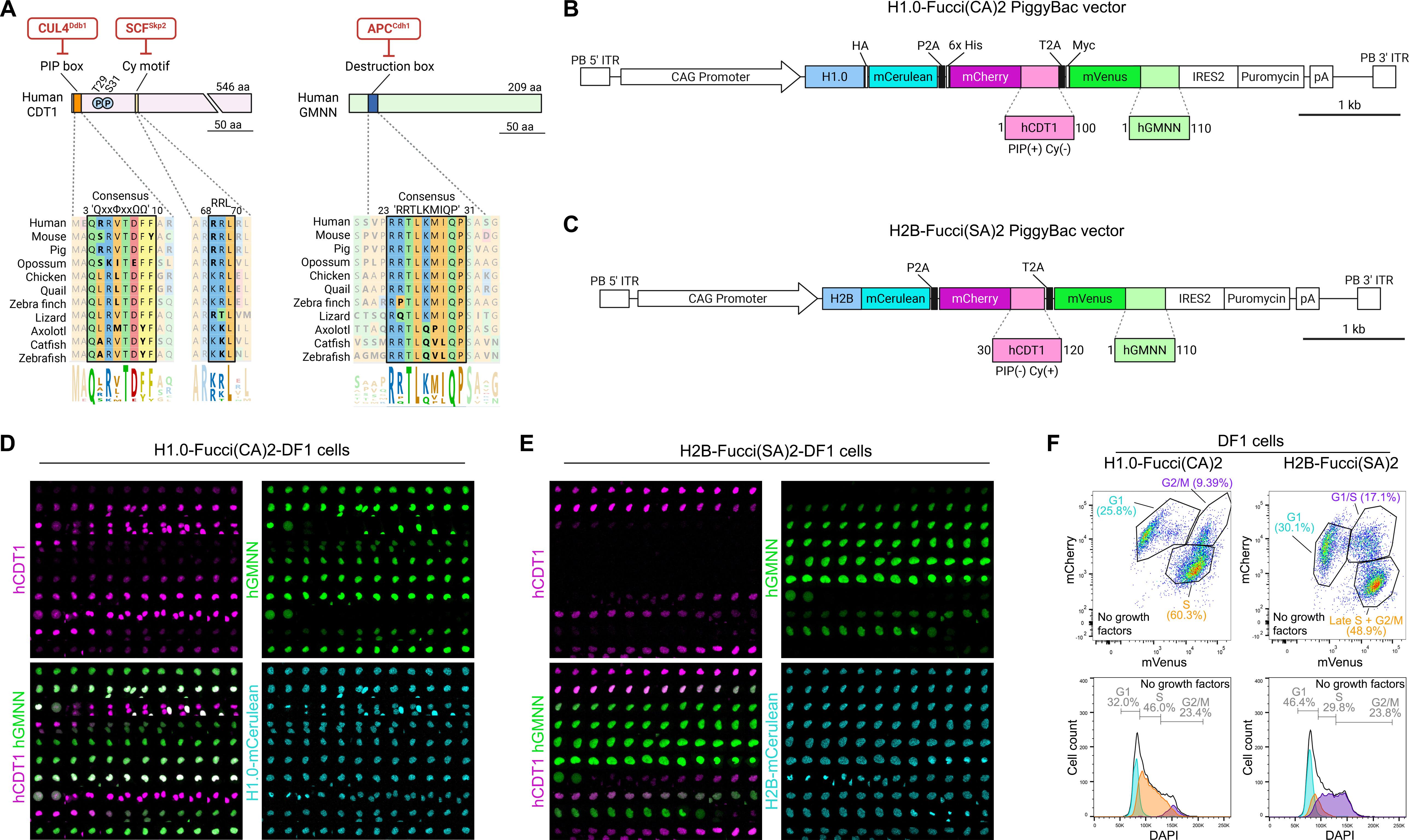

### Supplementary Figure 2

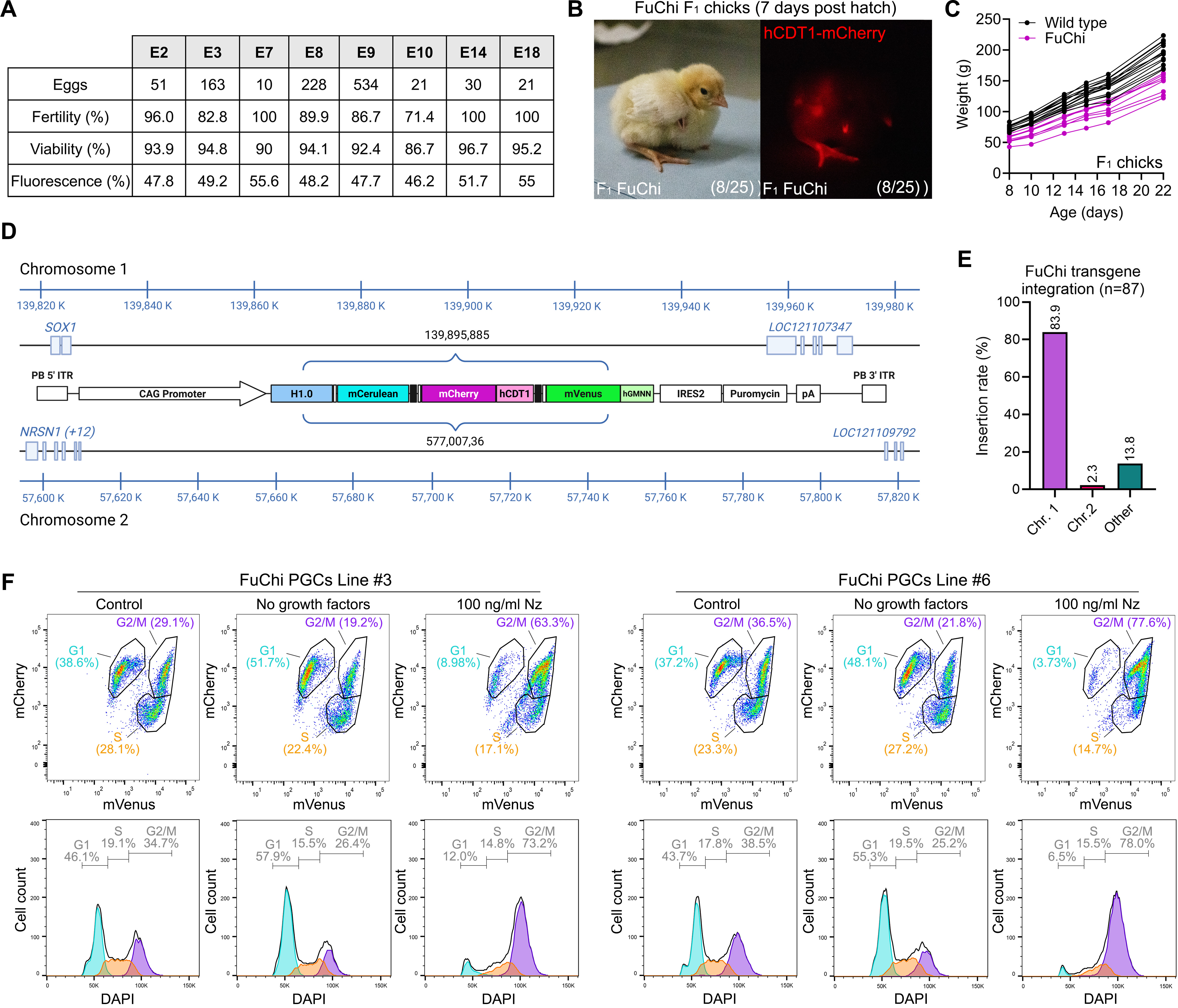

### Supplementary Figure 3

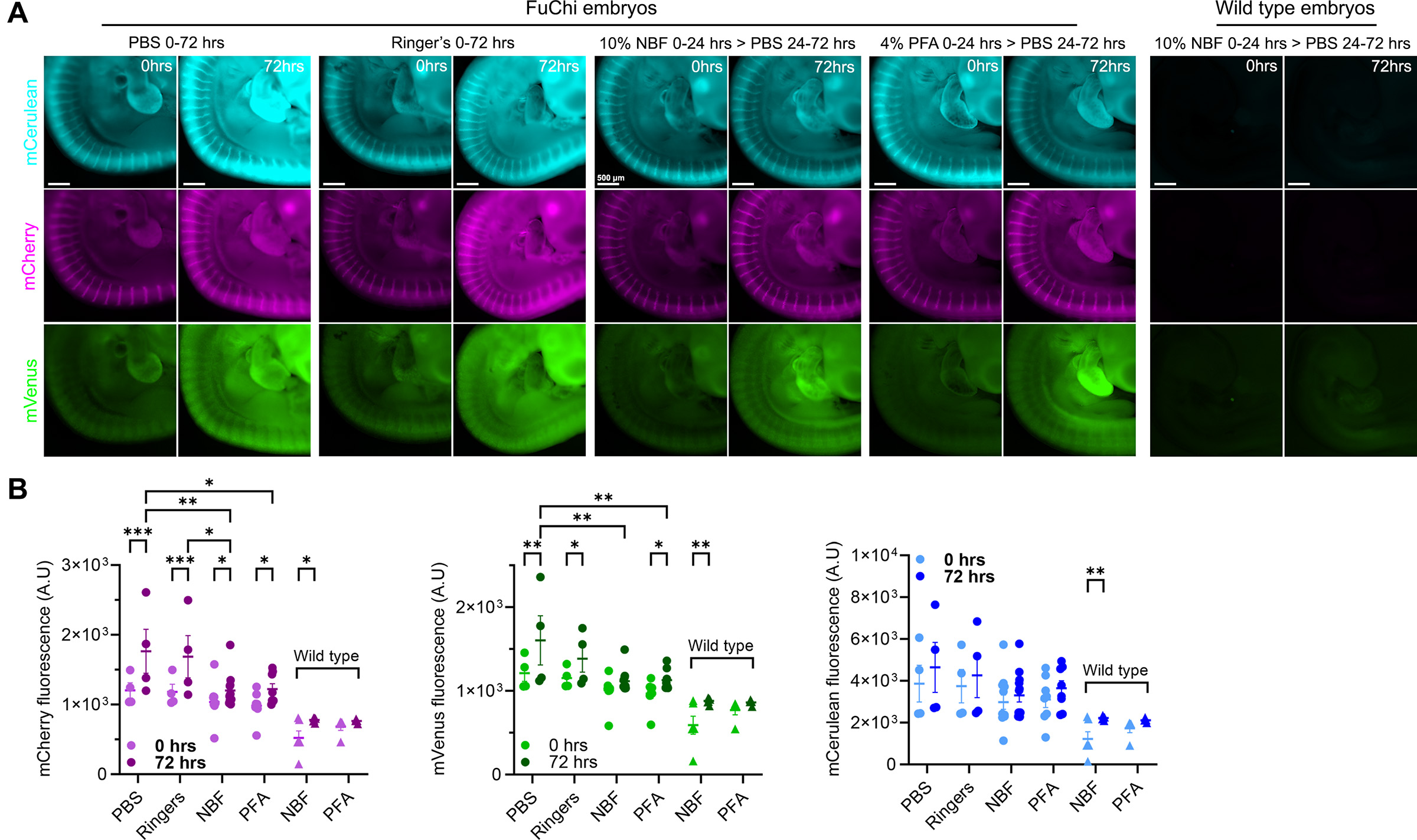

### Supplementary Figure 4

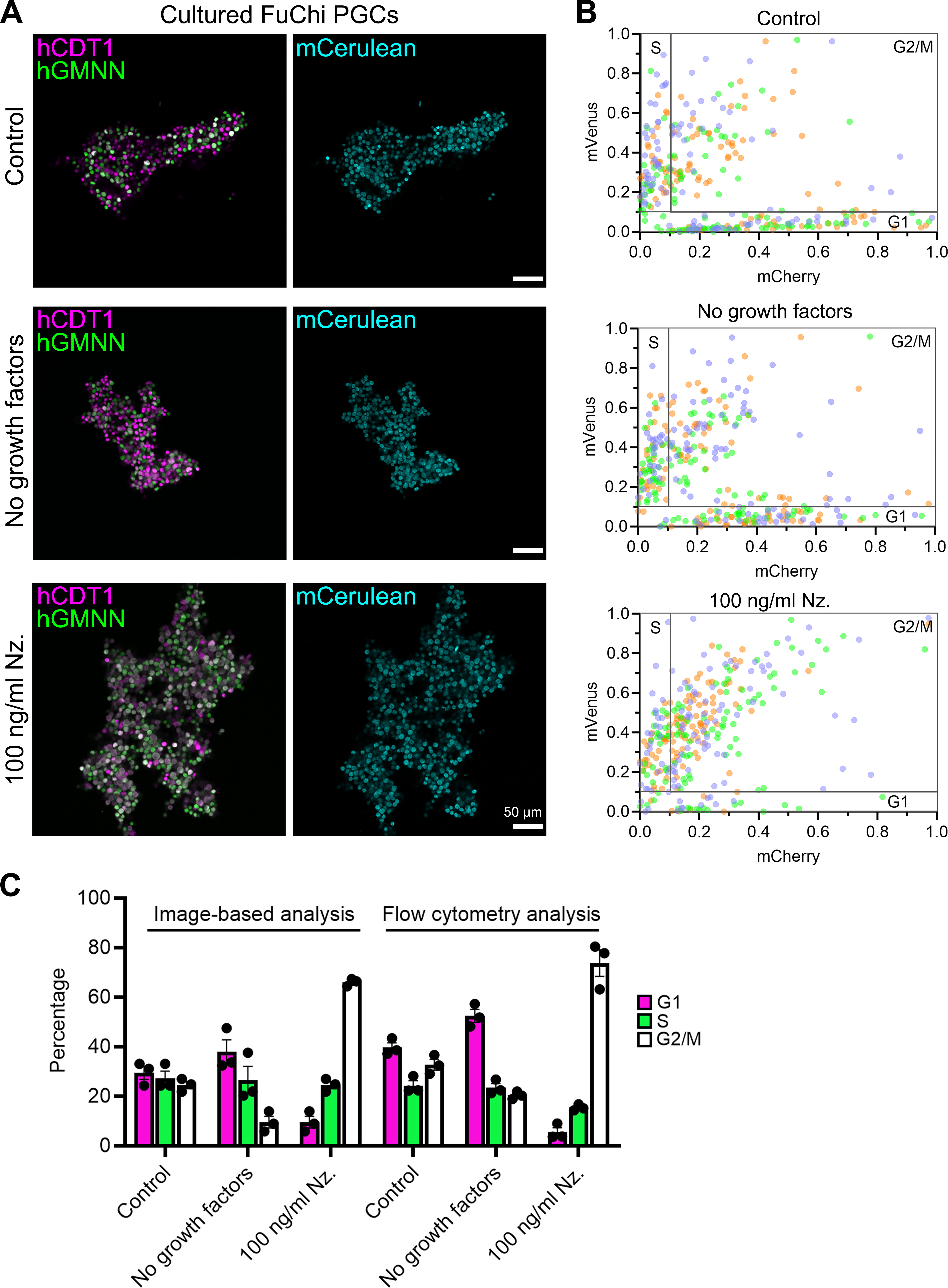

### Supplementary Figure 5

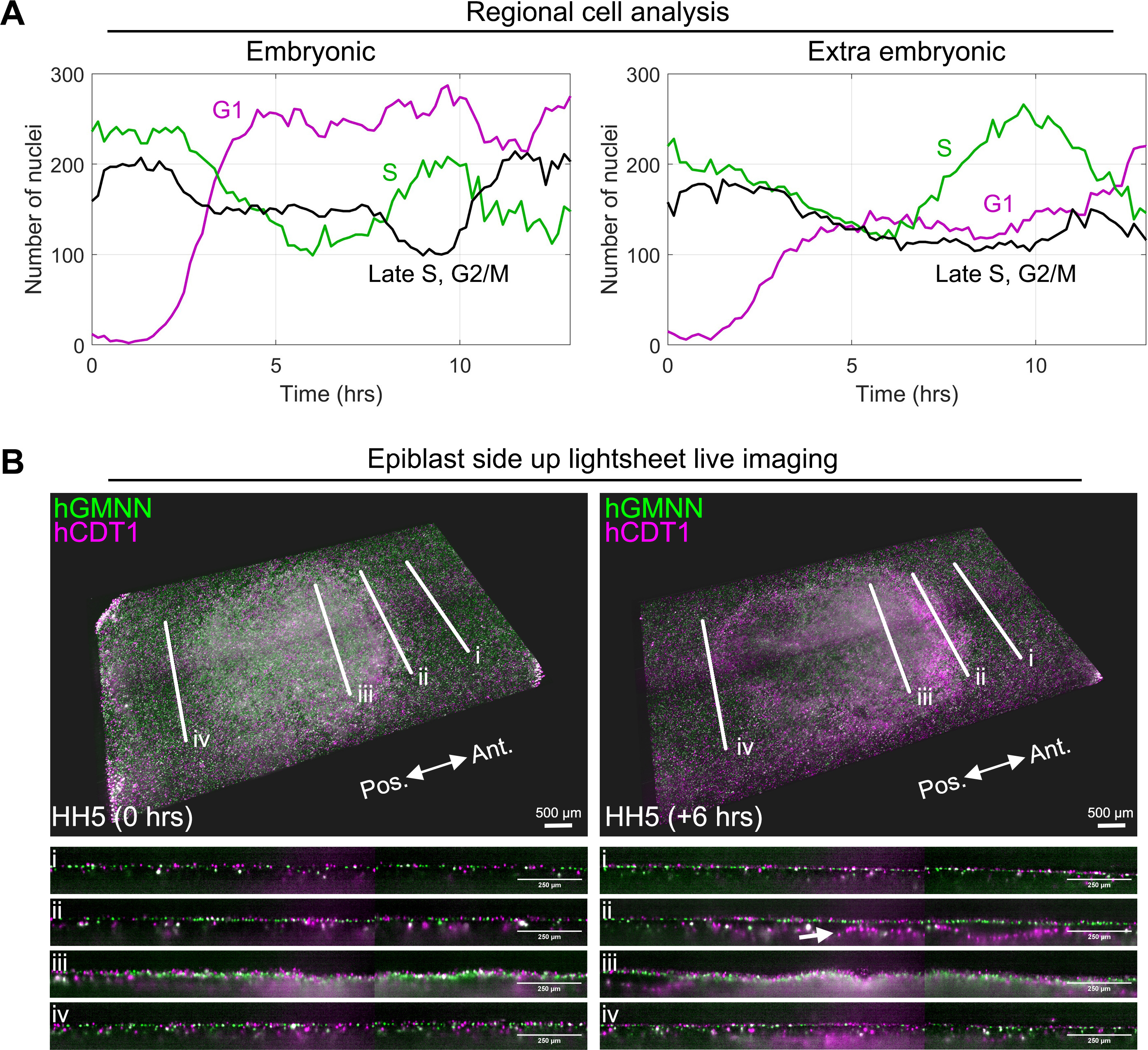
